# Cryo-EM map anisotropy can be attenuated by map post-processing and a new method for its estimation

**DOI:** 10.1101/2022.12.08.517920

**Authors:** Ruben Sanchez-Garcia, Guillaume Gaullier, Jose Manuel Cuadra-Troncoso, Javier Vargas

## Abstract

One of the most important challenges in cryogenic electron microscopy (cryo-EM) is the substantial number of samples that exhibit preferred orientations, which leads to an uneven coverage of the projection sphere. As a result, the overall quality of the reconstructed maps can be severely affected, as manifested by the presence of anisotropy in the map resolution. Several methods have been proposed to measure the directional resolution of maps in tandem with experimental protocols to address the problem of preferential orientations in cryo-EM. Following these works, in this manuscript we identified one potential limitation that may affect most of the existing methods and we proposed an alternative approach to evaluate the presence of preferential orientations in cryo-EM reconstructions. In addition, we also showed that some of the most recently proposed cryo-EM map post-processing algorithms can attenuate map anisotropy, thus offering alternative visualization opportunities for cases affected by moderate levels of preferential orientations.

## 1. Introduction

Ten years after the beginning of the resolution revolution [1], cryogenic electron microscopy (cryo-EM) has become one of the most popular approaches in structural biology, delivering thousands of high-resolution structures for a wide range of macromolecules. Despite being used routinely, cryo-EM is far from being a fully automatic technique and important challenges have yet to be resolved. One of such recurring problems is the preferred orientations that some specimens tend to exhibit when vitrified interfaces [2].

Preferred orientations are a consequence of the interactions between the particles and the air-water or substrate-water interfaces [3,4]. This may cause damage or degradation of the macromolecular structure, compositional heterogeneity and nonuniform distributions of angular projections leading to regions of the projection sphere with poor coverage [5]. As a result, the reconstructed maps may be artefactual because of partial to complete sample denaturation and/or tend to exhibit worse resolution in the direction parallel to the preferred orientation axis. For severe cases, important artefacts as map stretching could also appear [6]. These problems are generally described in terms of (directional) resolution anisotropy and can be explained by the lack of information generated by missing particle projections, as well as by the overabundance of certain projections that can hinder the reconstruction process [7].

It has been widely accepted that resolution anisotropy is mostly a manifestation of the preferential orientations problem, and given its ubiquity, several metrics based on this idea have been proposed. One of the first proposed metrics was the 3D-FSC, in which the gold standard FSC resolution estimation is computed in a directional manner by splitting the shells into cone sections perpendicular to the radius of the volume [6]. As a result, an independent FSC curve with an associated resolution estimation is obtained for each of the cones in which the sphere is discretized. As a summary of the per-cone resolution, the sphericity of the thresholded 3D-FSC volume was proposed. Based on a different principle, Naydenova and Russo [3] proposed the orientation distribution efficiency, which is a measurement of the sphericity of the point spread function derived from the distribution of angular assignments. More recent in [8,9] authors proposed a directional local resolution estimator based on the use of the monogenic signal over the EM map along different directions computed from directed cones.

All the methods presented above are based on analysing the directional resolution from a given 3D reconstruction [6,8,9] or a particle orientations distribution [3]. Consequently, the estimated directional resolution depends obviously on the particle orientation distribution. However, as it will be shown in the Results Section “Common map anisotropy metrics are affected by the shape of the specimen”, this magnitude also depends on the actual geometry of the sample to reconstruct, since the spectral signal to noise ratio (SSNR) at different directions may depend on the actual shape of the macromolecule. While this latter dependency has no impact when the metrics are used to compare different reconstructions of the same sample, it makes direct comparison of different specimens infeasible.

Map anisotropy metrics are invaluable diagnostic tools that can be employed to assess if a given approach can alleviate the preferred orientations problem. Examples of such experimental approaches are the usage of surfactants support films as a barrier [10], alternative types of grids [11–13], reducing the time that the sample dwells in the thin liquid film before vitrification [14] or re-imaging after tilting the sample to introduce missing views [6]. Although these previously mentioned approaches have been found to be useful, the presence of preferential orientations is still a major hurdle for cryo-EM as it limits the throughput of the technique and could generate artefacts leading to incorrect conclusions. From a computational perspective, although it is generally known that different algorithms exhibit different degrees of robustness against this problem, only little research has been carried out about this topic [7] and, to the best of our knowledge, only the Phenix anisotropic sharpening method was explicitly designed to perform anisotropy correction at the map level [15–17]. However, our experience leads us to think that some of the recently published map postprocessing algorithms could also alleviate the effect of preferred orientations in post-processed maps without being explicitly designed for that purpose. To the best of our knowledge, no formal study has been carried out to determine if such perception is accurate.

From a theoretical perspective, anisotropy reduction could be achieved by several computational mechanisms. First, anisotropy can be attenuated if the maps are isotropically filtered, or the frequencies are radially averaged in a way that homogenizes the information from different directions in the Fourier space. Consequently, the intensities in the directions that were originally well-represented in the anisotropic map are dampened whereas intensities from previously weak directions are boosted. Similarly, since high-resolution information is more affected by map anisotropy, attenuation can also be attained by low-pass filtering. These simple mechanisms can reduce the anisotropy of the map, making it more visually appealing at the cost of attenuating the high-resolution information. Alternatively, map anisotropy could also be reduced by including *a priori* information in the post-processing calculations so that the experimentally missing information is filled, at the risk of introducing potential extrapolation errors. Similarly, general hypotheses derived from *a priori* information to perform nonlinear modifications of the density could also “correct” the anisotropic properties of the map, but they may also introduce artefacts if the hypothesis do not hold true. While all the above strategies would result in a measurable reduction in map anisotropy, it is important to understand that the nature of the effect is different, and that the reduction may not necessarily translate in maps more similar to the ground truth (e.g., the atomic model). In the latter case, the anisotropy reduction could be considered as apparent and, although less desirable, it could be still beneficial for visualization purposes if the disagreement is not caused by artefacts. Serve as example a low-pass filtered version of a noisy map, which is easier to visualise at the cost of dampening high frequency information. On the other hand, approaches based on *a priori* information could in principle produce a genuine reduction of anisotropy depending on the accuracy of the prior. The crux of the matter is that there is currently no way to validate whether this inpainting of unobserved data accurately represents the actual underlying structure, and therefore adding missing information carries significant risks.

Even if it were only an apparent effect, and regardless about its desirability, map anisotropy reduction is already happening in many popular post-processing algorithms and the users should be fully aware of it, as it could result in easier visualization and interpretability of maps in some cases while potentially leading to problematic situations in some others. Therefore, we strongly recommend to the users to always validate their post-processed maps using the original, raw reconstructed map: if a feature is present in the post-processed map but not in the original one, it is safer to assume that it does not exist.

In this work, we illustrate a limitation that may affect all currently employed methods for map anisotropy estimation, and we propose an alternative approach to evaluate the presence of preferred orientation in single particle cryo-EM maps not affected by this limitation. In addition, we analyse how different map post-processing algorithms can alleviate the presence of anisotropic directional resolution in single particle cryo-EM maps affected by preferred orientations. Our results show that most of these approaches have a mitigating effect for cases exhibiting moderate levels of anisotropy.

## 2. Results and discussion

### 2.1. Common map anisotropy metrics are affected by the shape of the specimen

All the map anisotropy estimation methods described in the introduction are based on analysing the directional resolution from a given 3D reconstruction [6,8] or a particle orientations distribution [3]. The obtained directional resolution depends obviously on the particle orientation distribution. However, this magnitude also depends on the actual geometry of the sample to reconstruct. Note that the spectral signal to noise ratio (SSNR) at different directions may depend on the actual shape of the macromolecule. Thus, based on the relation between the FSC and SSNR [18], we can conclude that the 3D-FSC (or the efficiency) may also depend on the actual shape of the macromolecule. As example, consider a narrow and long filament along the z-axis. This structure will provide high Fourier amplitudes mostly localized close to the x-y plane and to the z axis. Consequently, in the presence of noise, the SSNR will show an anisotropic shape related with the macromolecular geometry. In Figure 1, we display some experimental cases where their SSNRs show clear anisotropy at medium-high resolutions related to the macromolecular geometry. To estimate the SSNR or the map confidence, we have computed in this figure the map normalized differences between corresponding half maps in Fourier space, which is given by 𝑀𝐷 = |𝑉_1_ − 𝑉_2_|⁄(|𝑉_1_| + |𝑉_2_|), with 𝑉_1_ and 𝑉_2_ the Fourier transforms of the half maps. Note that the smaller this magnitude at a given Fourier coordinate the higher the map confidence at this frequency. These cases comprise high-quality examples without and with low symmetry and correspond to cryo-EM maps with EMDB accession codes EMD-8910 (Figure 1a), EMD-29350 (Figure 1b) and EMD-13202 (Figure 1c). These maps have reported FSC- resolutions of 3.0, 2.5 and 3.1 Å respectively and map anisotropy is not visually present. In Figure 1, we also show for each map their corresponding slices of Fourier map amplitudes at 𝑧 = 0, 𝑦 = 0 and 𝑥 = 0 planes and black circles indicating a resolution of 4.0 Å. As can be seen from Figure 1, these maps clearly show strong signs of SSNR anisotropy at medium-high resolutions related with the macromolecular geometry.

**Figure 1.**
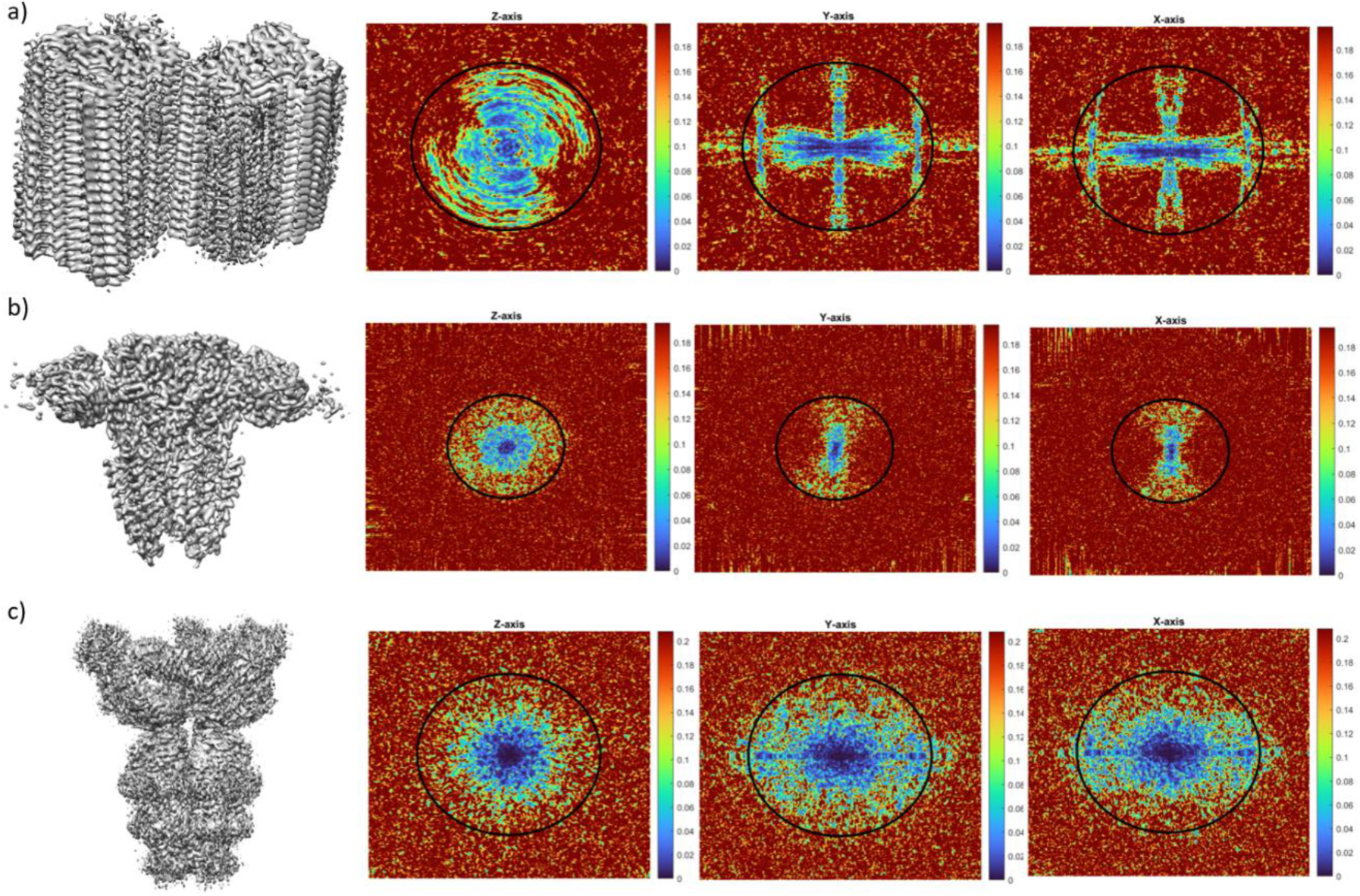
3D-FSC apparent anisotropy induced by the macromolecule shape. Each row shows an experimental cryo-EM map EMD-8910 (a), EMD-29350 (b) and EMD-13202 (c) and corresponding slices of Fourier amplitudes along 𝑧 = 0, 𝑦 = 0 and 𝑥 = 0 Fourier planes. The black circles indicate a resolution of 4.0 Å. The colorbar represents Fourier amplitude values at Fourier slices.

We also carried out tests in which we generated a simulated cryo-EM map from the atomic model of the influenza hemagglutinin (HA) trimer and we corrupted it by adding isotropic gaussian noise. Consequently, the map is perfectly isotropic and an ideal metric should not report any level of anisotropy. In Figure 2, we show the results obtained by the 3D-FSC [6] a), the efficiency [3] b), MonoDir [8] c-e) and our proposed approach f) for the simulated map. The computed sphericity for the 3D-FSC method was 0.67 and 0.85 at FSC thresholds 0.143 and 0.5 respectively. The reported efficiency calculated with the second technique was 0.81 with mean and standard deviation PSF resolution of 3.40 Å and 0.32 Å respectively. In Figure 2 c-e), we display the lowest, highest and half interquartile resolutions at some slices as computed by MonoDir, showing different values depending on the plane.

**Figure 2.**
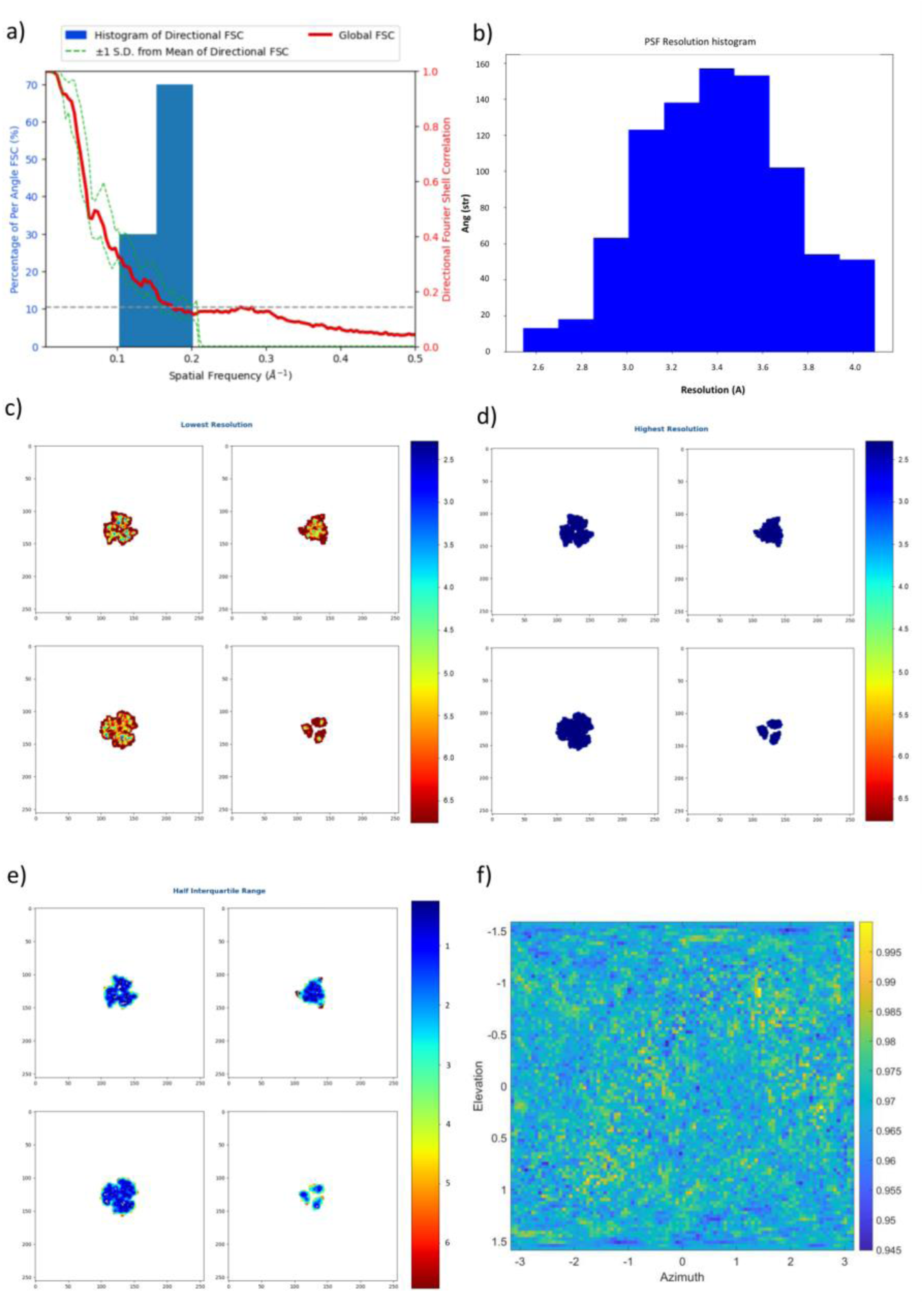
Results obtained for the simulation of the soluble portion of the small influenza hemagglutinin (HA) trimer by the 3D-FSC [6] a), the Efficiency [3] b), MonoDir [8] c-e) and the approach proposed in this manuscript f). Note that in f) the colorbar represents the noise power of the noise masked map along different directions given by the elevation and azimuth angles and normalized by the value at the direction providing maximum noise power. Contrary to what it is expected for a perfectly isotropic map, the 3D-FSC per-cone standard deviation a) is not zero, the histogram b) of resolution per angle is not narrow and the different planes of local-directional resolution computed with MonoDir are not at the same values. On the contrary, our approach shows almost constant values near 0.95. Consequently, all the studied methods except our proposed approach consider that the resolution of the ideal simulated volume is anisotropic.

As can be concluded from these results, all methods except our proposed approach evaluated a perfectly isotropic simulated map as anisotropic. This limitation, far from being exclusively theoretical, may have an important consequence as anisotropy estimation across different specimens are not comparable. Note however, that these metrics are perfectly valid to analyse the anisotropy of different reconstructions of the same sample since they share the same shape.

### 2.2. New anisotropy method results

We developed a new method to estimate the map anisotropy based on the directional distribution of noise power (please see section 3.3 New anisotropy method). Figure 2 f) shows the results obtained by our approach on the same simulated, isotropic density map as in the previous section. As it can be concluded from direct inspection of the 2D heatmap, the density is evaluated as isotropic because the noise power, which is close to 1, does not change significantly along different directions.

We have also evaluated our approach on the experimental maps of the influenza hemagglutinin (HA) trimer. Figure 3 a) and b) display the results provided by our method for the reconstruction using untilted micrographs and the reconstruction using micrographs tilted at 40° respectively. For the untilted case, the 2D heatmap shows high values for elevations between −1 to 1 rad approximately (note that an elevation of 0 rad corresponds to side views), while for the 40° tilted case, this range is reduced to elevations between −0.5 to 0.5 rad. This is in agreement with the better quality of the map with tilted particles. Also note that the 2D heatmap corresponding to the 40° case shows high values at elevations close to −1.5 and 1.5 rad that indicate lower levels of information for those elevations. However, since these elevations correspond to top views (particles with tilts close to 0°), which do not introduce much information in the reconstruction, their impact in the overall quality is minor. Moreover, for the untilted case (Figure 3 a)), three clear preferred orientations within the asymmetric unit can be found for elevation values close to zero (side views) that cannot be observed for the 40° case (Figure 3 b)), which likely correspond to particles incorrectly aligned by the angular refinement approach.

**Figure 3.**
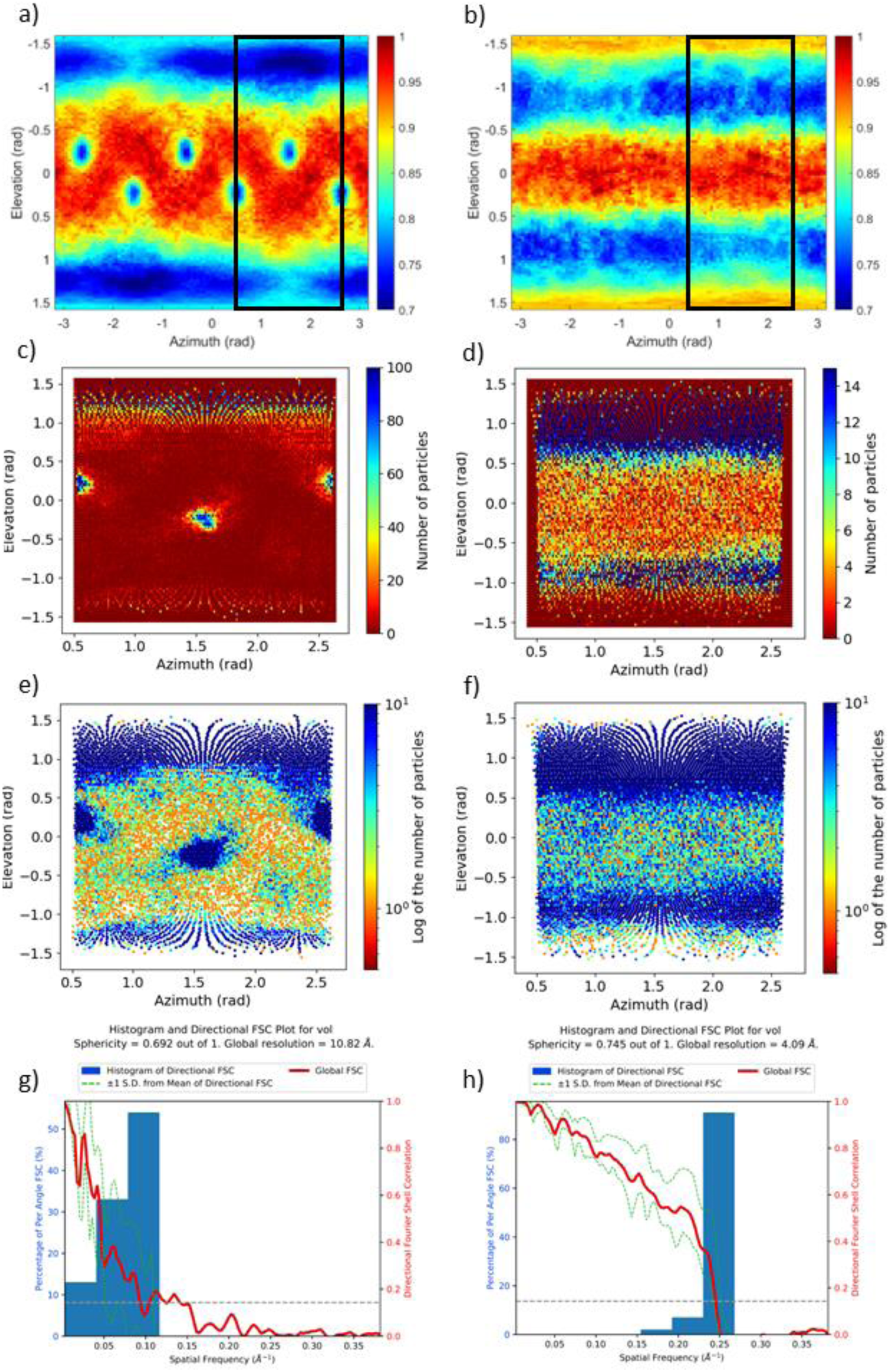
Results obtained by the proposed method for experimental maps obtained from untilted (a) and 40° tilted single particles of the influenza hemagglutinin (HA) trimer (b). The particle angular distribution as obtained by Relion is shown in c) as counts per orientation or as the logarithm of the counts in e), as reflected in the colorbar y. The same plots are shown in d) and f) for the tilted case. The colorbars in a) and b) represent the noise power of the noise masked map along different directions given by the elevation and azimuth angles and normalized by the value at the direction providing maximum noise power. The elevations and azimuths are given in radians. The results obtained by the 3D FSC method are shown in (g) and (h) for the untilted and 40° tilted cases respectively.

In Figure 3 c) and d) we include the corresponding particle angular distribution obtained from the particle’s orientation computed by Relion [19,20]. Figures 3 e) and f) show the logarithm of the corresponding angular distributions. Note that the Y-axes in these figures are inverted with respect to the ones in Figure 3 a) and b) and that the elevation angle used by our approach has its origin in the equatorial plane of the projection sphere, while the tilt angle used by Relion has its origin in the north pole. As can be seen from these figures, the 2D maps obtained by our approach and from the particle distributions are very similar, including the three clear preferred orientations for elevation values close to zero (side views) in the 0° case, which cannot be observed in the 40° case. Furthermore, Figure 3 g) and h) show results obtained using the 3D-FSC approach. The Sphericity and Global resolution calculated from the map-model 3D-FSC are 0.692 and 10.82 Å for the 0° tilt and 0.745 and 4.09 Å for the 40° tilt, respectively. These results indicate that the 3D-FSC performance is significantly lower for the 0° case, likely due to many particles being incorrectly aligned, despite the existence of a pronounced preferred orientation issue at high elevation values in both cases. This outcome suggests that while the particle orientation distribution (or the noise power distribution provided by our method) is effective in identifying preferred orientation issues, it cannot directly assess the impact of these issues on map quality. Conversely, complementary metrics such as the 3D-FSC can offer insights into the effect of these issues on map quality, especially in cases where the shape effect of the macromolecule can be disregarded. Therefore, both metrics should be considered complementary.

We also evaluated our newly proposed method on β-galactosidase maps in which anisotropy was artificially induced by removing 95% of the particles assigned to tilt angles above 80° or 70° (Relion convention, side views). The results obtained by the proposed approach are shown in the first row of Figure 4. As can be seen from this figure, the case where no particle was removed (0% removed) shows low values at two orientations corresponding to side views approximately (elevation close to 0 rad). Oppositely, the cases with induced anisotropy by removing particles at tilt angles >80° or >70° show high values at elevation close to 0 rad, revealing the lack of side view (or close-to-side view) particles in these cases. Notice also that the case with particles removed at tilt angles >80° exhibit lower values for small elevation angles between −0.5 and 0.5 rad (high tilt angles) than the case with particles removed at tilt angles >70° showing that in the latter case there are less side view particles. The black rectangles in the figures correspond to the regions shown in the 2D particle angular distributions displayed in the second and third rows of Figure 4, which are in line with the results obtained by our proposed approach.

**Figure 4.**
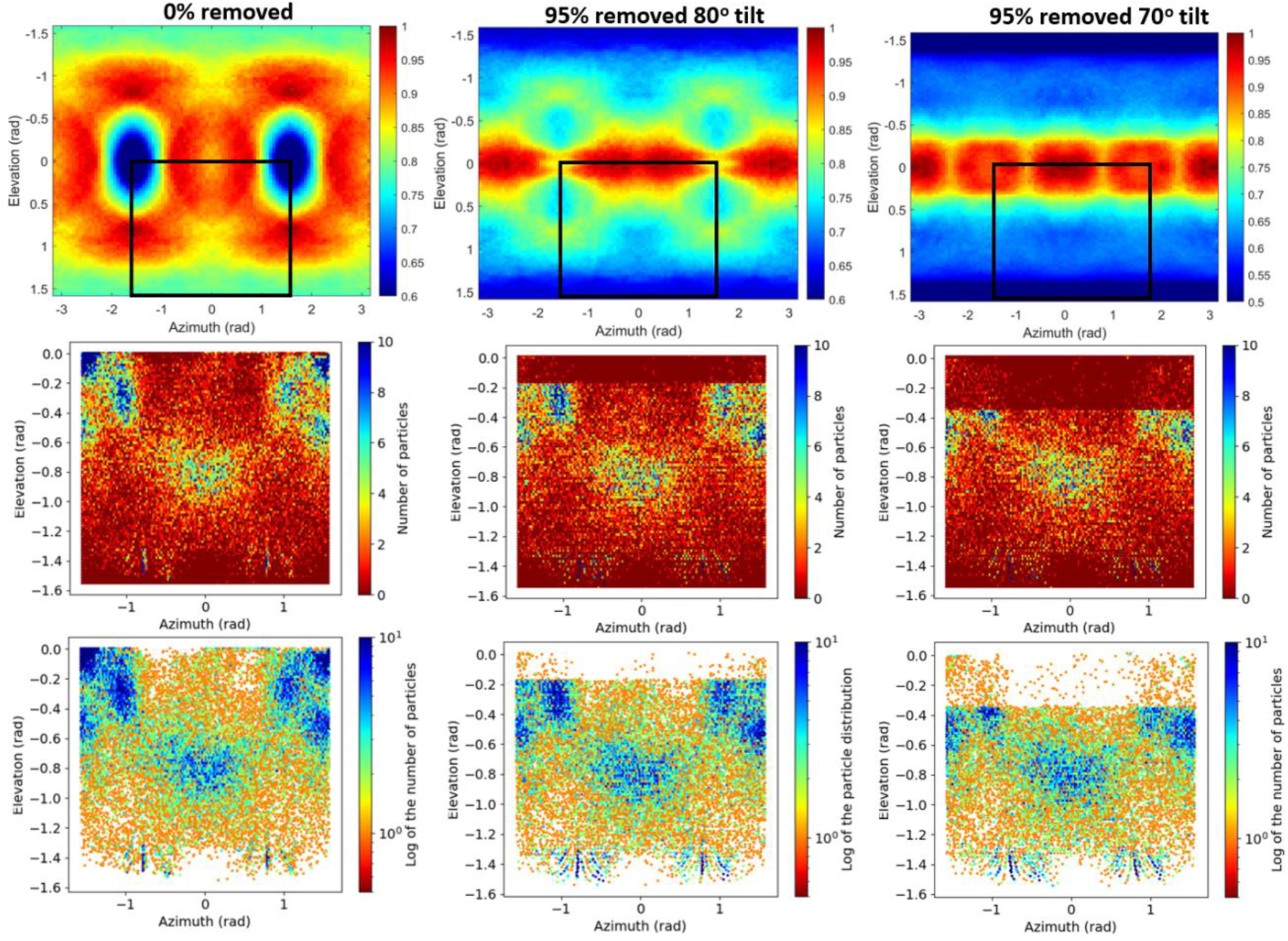
Results obtained for reconstructions of the β-galactosidase complex (EMPIAR-10061) consisting of a standard reconstruction of the β-galactosidase at 3.15 Å and two reconstructions in which anisotropy was artificially induced by removing 95% of the particles assigned to tilt angles above 80° or 70° (side views). The first row shows the results obtained by the proposed method. The black rectangles in the figures correspond to the regions shown in the 2D particle angular distributions displayed in the second (particle count) and third (logarithm of the particle count) rows. The colorbars represent the noise power of the noise masked map along different directions given by the elevation and azimuth angles and normalized by the value at the direction providing maximum noise power.

### 2.3. Nonlinear post-processing methods can attenuate map anisotropy

In order to address the question of whether map post-processing can attenuate map anisotropy, we computed the 3D-FSC [6] of post-processed maps using DeepEMhancer [21], LocScale2 [22,23], LocSpiral [24], LocalDeblur [25], and Phenix anisotropic sharpening [15] against the maps simulated from the corresponding atomic models at the FSC global resolution for three different datasets. Note that we have not used our proposed anisotropy method to evaluate the post-processed maps, since some of the sharpening methods used perform automatic masking that precludes the use of our approach. However, as previously explained, the 3D-FSC is perfectly valid for comparing different maps of the same macromolecule.

The first dataset is composed of three different versions of the β-galactosidase complex (EMPIAR id 10061, [26]), a standard reconstruction at 3.15 Å and two reconstructions in which we artificially introduced preferred orientations by removing 95% of the particles assigned to tilt angles above 80° or 70° (side views). Figure 5 top displays the standard deviation of the per-cone FSC computed on the different maps against the corresponding atomic model. As expected, the standard deviation for all maps and frequencies is small when we consider the case not affected by induced anisotropies (0% removed), indicating that the maps are mostly isotropic in this case. Equally expected, the standard deviation of the per-cone FSC for the reconstructed maps (not postprocessed, blue curves) increase when we severely reduced the number of particles at high tilts, especially in the high-resolution range. For example, at the FSC global resolution frequency (marked in the figures with a dashed vertical line), the FSC standard deviation increases from <0.04 for the reconstructed map (blue curve) for the case (0% removed) to values above 0.05 and 0.08 when 95% of the particles with tilt angles >80° and tilt angle >70° are removed respectively.

**Figure 5.**
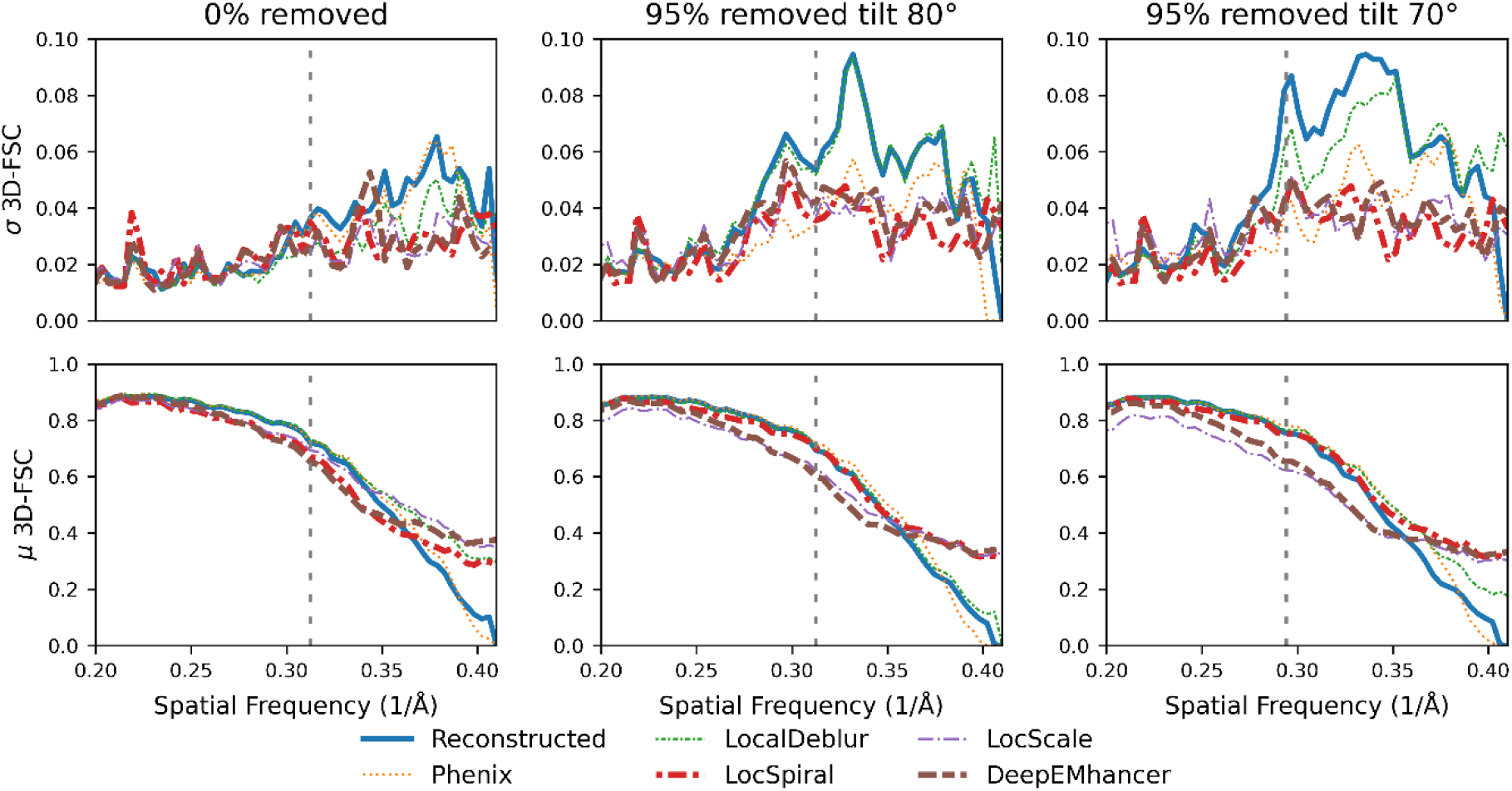
Per-cone FSC (map to model) standard deviation (top) and mean (bottom) calculated with 3D-FSC for three different versions of the β-galactosidase complex: reconstructed with all particles (left) and reconstructed with 95% of the particles with tilt angle >80° (middle) and >70° (right) removed. The blue line represents the reconstructed map whereas the brown, purple, red, orange, and green lines represent the maps obtained when DeepEMhancer [21], LocScale2 [22,23], LocSpiral [24], Phenix [15] and LocalDeblur [25] are employed on the reconstructed map. The FSC global resolution is displayed as a dotted vertical line. See Supplementary Figure S3 for an alternative representation of the data.

From direct observation of the standard deviation for the different post-processed maps for the >80° and the >70° cases, we can conclude that all post-processing algorithms are able to reduce the standard deviation in the high-resolution region, thus apparently, they can reduce map anisotropy for these cases. While the effect of the LocalDeblur algorithm is quite small or even negligible in the high-resolution frequencies, the other post-processing methods are able to reduce the standard deviation of the per-cone resolution to levels comparable to the experimental reconstruction.

Moreover, as Figure 5 bottom shows, for several of the studied methods, the average per-cone FSC at high frequencies is better than for the reconstructed map (blue), especially when the anisotropy is more severe. The best performing algorithms for this example showed similar or better mean per-cone FSC with much smaller standard deviation than the reconstructed map, which confirms that the post-processing methods not only could attenuate the map anisotropy, but also that some of them generated maps more similar to the ground truth.

The second example is the integrin αvβ8 in complex with L-TGF-β (EMD-20794, [27]). This map is a 3.3 Å reconstruction with a poorly defined region in which anisotropy is noticeable. As in the previous example, Figure 6 top displays the standard deviation of the per-cone FSC computed on the different maps against the corresponding atomic model. As it can be observed from this figure, the per-cone standard deviation of the experimental map (not postprocessed, blue curve) is quite high at frequencies close to the global resolution (marked with a vertical dashed line), with higher levels than in the previous experiment. Despite that, and in the same manner as in the previous example, all post-processing methods except LocalDeblur are able to reduce the anisotropy of the map as measured by the reduction in per-cone FSC standard deviation at high frequencies, with DeepEMhancer and LocSpiral exhibiting the strongest reductions. Figure 6 bottom displays the average per-cone FSC computed between the different maps and the atomic model. This figure shows that LocScale and Phenix sharpening obtained higher FSC values than the experimental map for all frequencies, while the other methods only show better FSC in the high frequency range. Selected regions of the resultant maps are displayed in Figure 7, where it can be visually confirmed that post-processing methods offered more detailed maps.

**Figure 6.**
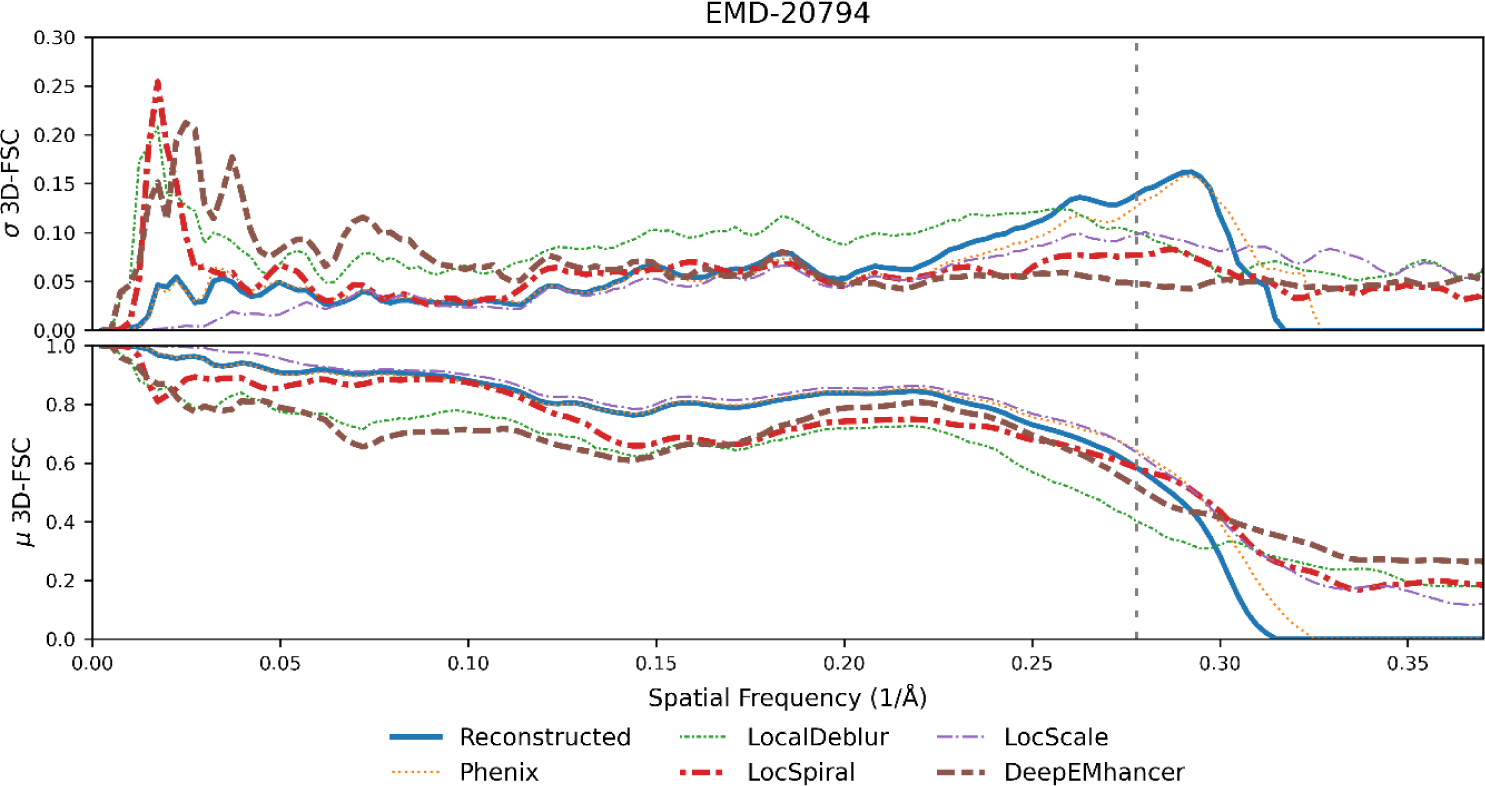
Per-cone FSC standard deviation (top) and mean (bottom) calculated with 3D-FSC for the EMD-20794. The blue line represents the reconstructed map whereas the brown, purple, red, orange, and green lines represent the maps obtained when DeepEMhancer [21], LocScale2 [22,23], LocSpiral [24], Phenix [15] and LocalDeblur [25] are employed on the reconstructed map. The FSC global resolution is displayed as a dotted vertical line. See Supplementary Figure S4 for an alternative representation of the data.

**Figure 7.**
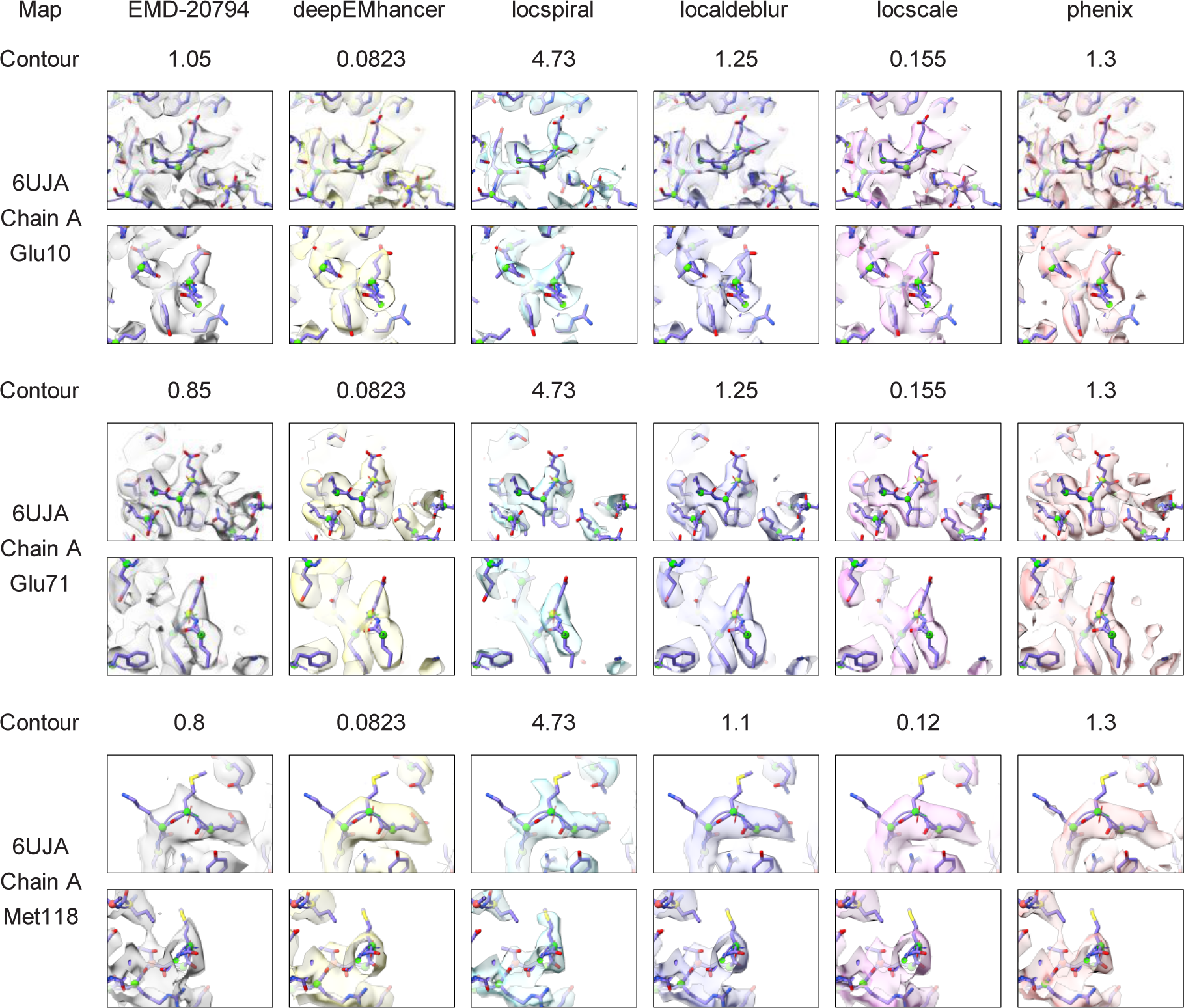
Reconstructed map (grey) for the EMD-20794 and the results of applying DeepEMhancer (yellow, [21]), LocSpiral (cyan, [24]), LocalDeblur (purple, [25]), LocScale2 (pink, [22,23]) and Phenix anisotropic sharpening (salmon, [15]) to the reconstructed map. The atomic model 6UJA is shown as reference. Three different locations viewed from two different orientations are displayed. They correspond to chain A residues 10, 71 and 118 and are displayed. Contour levels were manually selected to maximize the level of side chain inclusion.

The last example is the influenza hemagglutinin trimer reconstructed by Zi Tan et al. in their seminal work about how to address the preferential orientation problem [6]. We used both the map reconstructed from untilted micrographs (EMPIAR ID 10096), and the map reconstructed with micrographs tilted 40° (EMPIAR ID 10097). While the former is an exemplar of a specimen severely affected by the preferential orientation problem, the latter is a corrected version in which although still present, the map anisotropy was attenuated.

When we repeated the same experiment as in the other examples with the untilted hemagglutinin (see Figure 8), the results showed that none of the post-processing methods was able to reduce the standard deviation of the per-cone FSC computed from the experimental map (not postprocessed, blue curve). Indeed, some of them made the situation even worse (see Supplementary Figure S2), which is in line with what Sorzano et al. observed [7]. However, when the post-processing methods were employed on the map reconstructed on the tilted micrographs, in which the standard deviation decreased by a factor 2 compared to the untilted case, we were able to observe again a reduction in the standard deviation, especially remarkable for DeepEMhancer. As in previous experiments, we also observed that many of the post-processing algorithms obtained better than experimental mean per-cone FSCs (Figure 8, bottom), which translate into maps that are more similar to the simulated map computed from the atomic model than the experimental one, as Figure 9 suggests.

**Figure 8.**
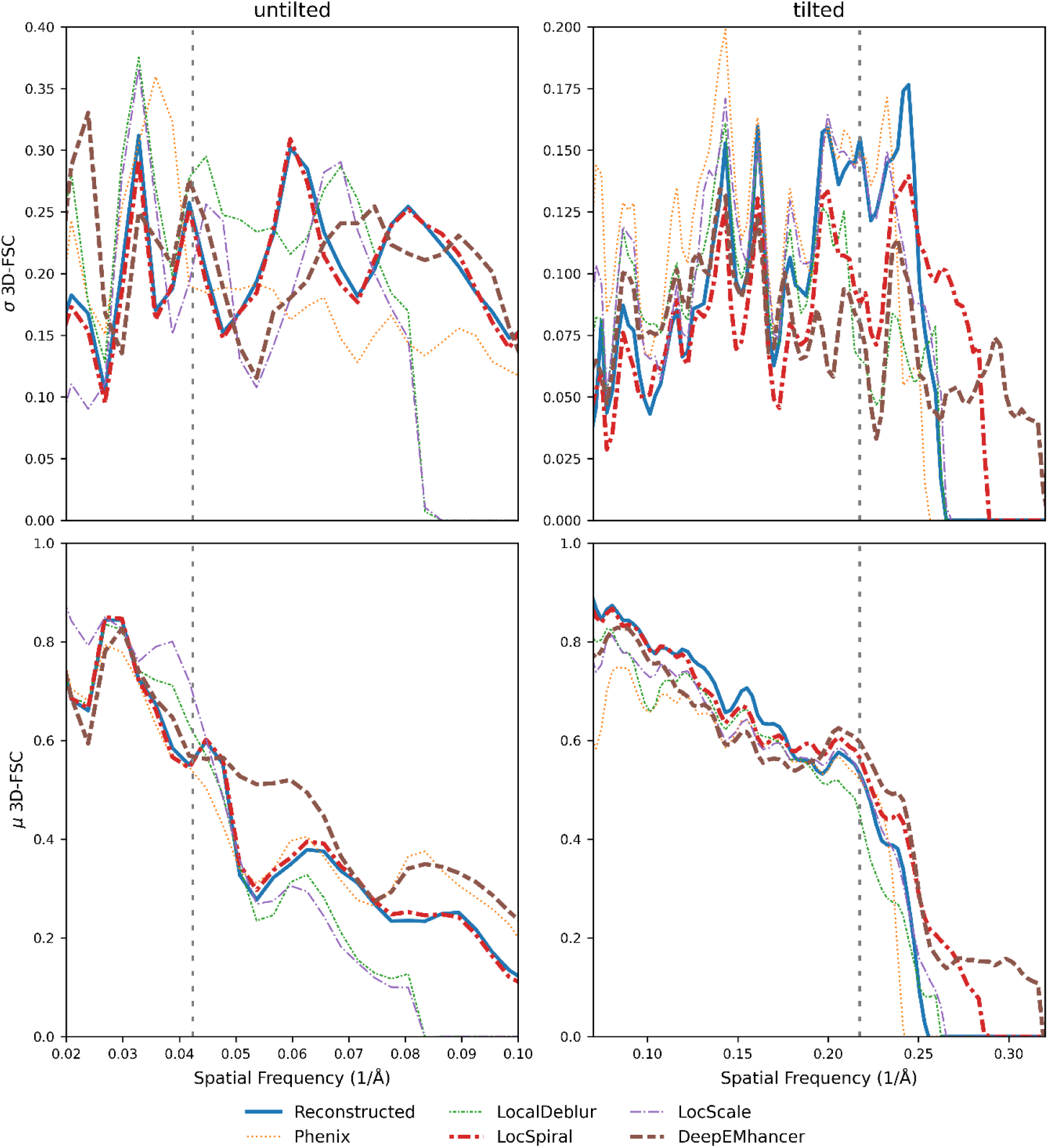
Per-cone FSC (map to model) standard deviation (top) and mean (bottom) calculated with 3D-FSC for two different versions of the influenza hemagglutinin trimer (HA): reconstructed with untilted micrographs (severe anisotropy problems) and reconstructed with micrographs tilted 40°. The blue line represents the reconstructed map whereas the brown, purple, red, orange, and green lines represent the standard deviation for the maps obtained when DeepEMhancer [21], LocScale2 [22,23], LocSpiral [24], Phenix [15] and LocalDeblur [25] are employed on the reconstructed map. The FSC global resolution is displayed as a dotted vertical line. See Supplementary Figure S5 for an alternative representation of the data.

**Figure 9.**
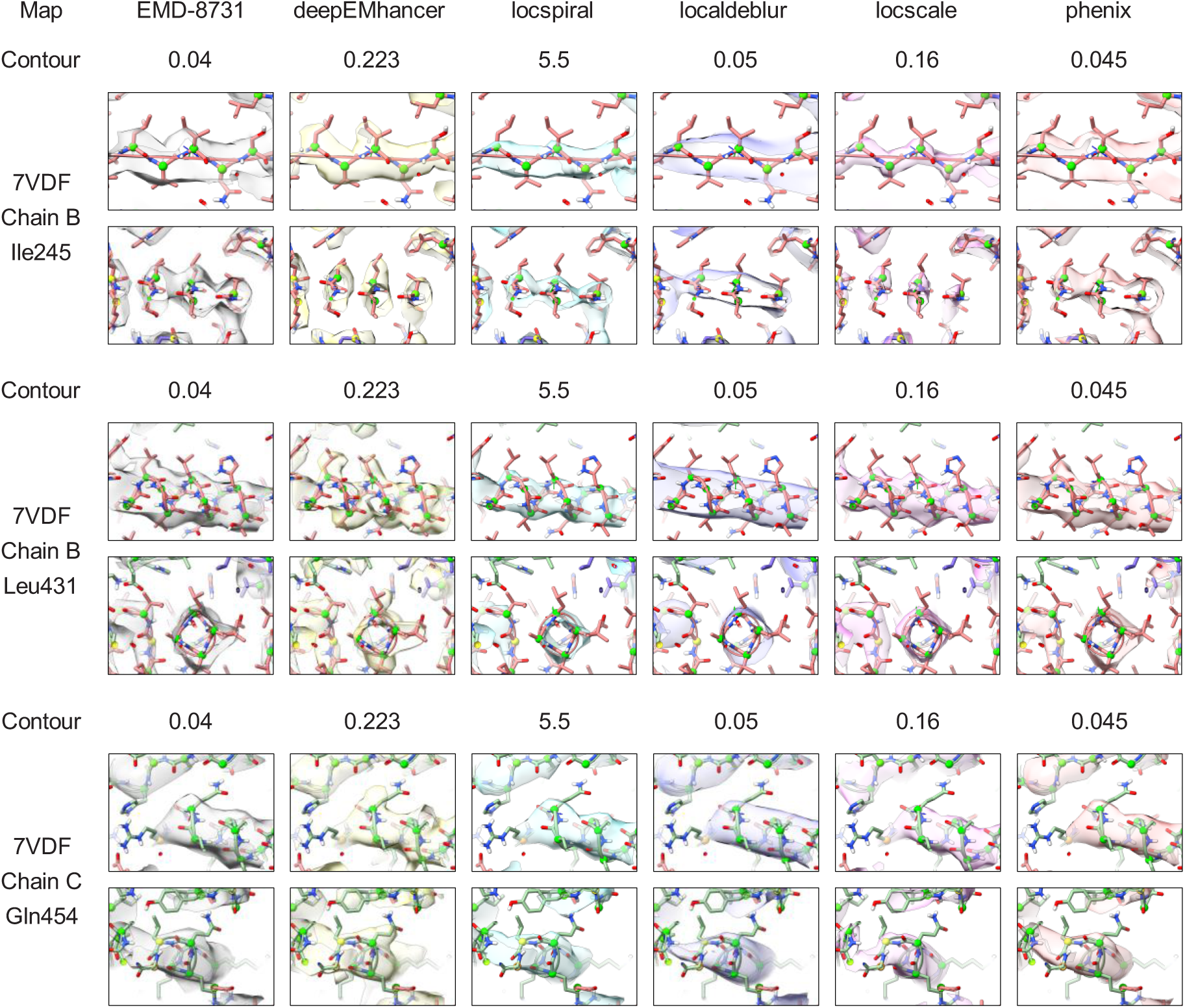
Influenza hemagglutinin trimer (HA) map reconstructed from tilted micrographs (grey), and the results of applying DeepEMhancer (yellow, [21]), LocSpiral (cyan, [24]), LocalDeblur (purple, [25]), LocScale2 (pink, [22,23]) and Phenix anisotropic sharpening (salmon, [15]) to the reconstructed map. The atomic model 7VDF is shown as reference. Three different locations viewed from two different orientations are displayed. They correspond to chain B residues 245 and 431, and chain C 454 are displayed. Contour levels were manually selected to maximize the level of side chain inclusion.

It could be argued that the simulated atomic densities that are used as ground truth and in some cases as source of priors are generated using isotropic atomic B factors. This means that each atom is modelled by an isotropic gaussian function with a given width, which introduces an incorrect bias since atoms are not anisotropic. As a consequence, some of the observed improvements obtained by the postprocessing approaches could be just a consequence of this assumed isotropy. Observe, however, that this assumption will only have a significant effect in the computed FSCs at high resolutions (∼3 Å or better), while our analysis is more focused on a range between 3-7 Å. Note that the resolutions at which this prior knowledge has a significant effect on the FSC curves can be observed at high-resolutions where the averaged FSC of the unsharpened map (blue curve) reaches zero, but the corresponding curves of most of postprocessed maps do not. Moreover, observe that all postprocessed maps have been masked with the same low-pass filtered mask. Thus, postprocessing methods introducing a priori information in the reconstructions (DeepEMhancer, LocSpiral, LocalDeblur and LocScale) produce average FSC curves that that do not reach zero at high resolutions (>3 Å), while the Phenix approach, which is based on B-factor correction not introducing a priori information, reaches zero at high resolutions. As explained in the DeepEMhancer publication [21], this behaviour is due to the *a priori* information introduced in the postprocessed reconstructions such as its isotropy at high resolutions, which can be observed in the FSCs for resolutions of about ∼3 Å or better. Therefore, this possible effect should not affect our analysis.

From these three experiments, we can confirm the capability of the map post-processing methods to apparently attenuate moderate amounts of map anisotropy, a conclusion also supported by some examples discussed on Twitter [28]. It can also be appreciated that the anisotropy attenuation effect is also dependent on the amount of anisotropy of the original map. Thus, for the highly anisotropic untilted hemagglutinin map, no reduction in anisotropy was measured, whereas for the tilted hemagglutinin and the EMD-20974 maps, the standard deviation of the directional resolution decreased from values around 0.15 to 0.07 at frequencies close to the global resolution while for the β-galactosidase with artificial preferred orientations, a less anisotropic map, the values were reduced from 0.09 to 0.03. For these three experiments, DeepEMhancer was the technique that exhibited the best attenuation capabilities in terms of per-cone FSC, while other techniques achieved better results in some specimens than in others. We have also shown that, depending on the specimen, different methods were able to improve the agreement (mean per-cone FSC) between the post-processed maps and the atomic model when compared to the experimental map.

These results should not be surprising, since, as stated in the introduction, the different selected post-processing methods are of different nature, thus their anisotropy attenuation and enhancement capabilities are different. Thus, in Phenix anisotropic sharpening, the only method explicitly designed to deal with anisotropy, the anisotropy of the maps is estimated and then employed to calculate isotropic scaling factors. In the other algorithms, anisotropy is not explicitly considered, but the different methods incorporate priors about the macromolecules or perform some calculations such as radial averaging or isotropic filtering that lead to more isotropic results.

DeepEMhancer is a representative of the first strategy, since it was trained using isotropic target maps and ideally, it should transform any input data, regardless of its amount of anisotropy, to isotropic data. Considering that DeepEMhancer can freely modify the phases and amplitudes of the experimental maps to make them more similar to what it considers the ideal map, it can potentially approximate the missing experimental information. These approximations could lead to an actual correction of the map anisotropy if the predictions are accurate enough, something that is not possible to know ahead, thus the risk of misinterpretation.

The second strategy, that is employed in LocalDeblur and LocSpiral, produces (locally) directionally uniform amplitudes that are an average of directional amplitudes. Particularly, in LocalDeblur, the maps are deconvolved with a symmetric gaussian kernel proportional to the local resolution estimated on the map, whereas in LocSpiral, the Fourier amplitudes are radially normalized. This reinforces previously weak directions in the anisotropic map leading to real space maps that appear smoother in all directions.

Finally, in LocScale, the amplitudes of the reference atomic model are used to rescale the amplitudes of the post-processed map, potentially filling missing experimental information if the atomic model is accurate enough. In addition, since the scaling step is conducted by comparing the radial averages of the atomic map and the experimental map, the result will be directionally uniform, as in the second strategy methods.

To visualize the effect of the different postprocessing approaches, we show in Figure S6 the logarithm of Fourier amplitudes for the tilted influenza hemagglutinin trimer reconstructions at slices *z* = 0, *y* = 0 and *x* = 0. As can be seen from this figure the different postprocessing methods, except for the case of LocalDeblur, tend to make the Fourier amplitudes more isotropic, which is in line with the results presented in Figures 5, 6 and 8.

Since all different methods can offer results of different quality, and it is not possible to tell which is best if no ground truth information is available, we would like to encourage the users to apply as many possible algorithms and compare and contrast their results, as the consensus of orthogonal methods is always more robust than the individual methods.

## 3. Materials & Methods

### 3.1. Datasets

#### 3.1.1. Artificial anisotropy

We processed the EMPIAR-10061 dataset [26] with Relion 3 [20] to obtain a β-galactosidase map at 3.15 Å (FSC unmasked) from 41,122 particles without applying any masking or post-processing operation. This map shows negligible amounts of anisotropy. Then, we removed 95% of the particles assigned to tilt angles above 70° degrees (Relion convention) producing a particle set of 26,409 particles which was again processed using Relion to obtain a map at 3.40 Å resolution. Finally, we additionally reconstructed another map after removing 95% of the particles with tilt angles above 80° degrees (32,941 particles in total) resulting in a map at 3.19 Å. This last map exhibit anisotropy levels between the raw map and the one obtained removing the particles at tilt angles > 70°. See Supplementary Figure S1 for a visual of the densities and the 3D-FSC plots.

#### 3.1.2. Experimental maps

The first map was the EMD-20794 [29], a 3.3 Å resolution reconstruction of the Integrin αvβ8 in complex with L-TGF-β which exhibits moderate levels of anisotropy. In our analysis. The associated atomic model that we employed for 3D-FSC calculations was the PDB ID 6UJA.

The second specimen was the influenza hemagglutinin (HA) trimer obtained by Tan et al. [6]. We employed the untilted version, severely affected by preferred orientations (EMPIAR ID 10096), and the tilted version (EMD-8731, EMPIAR ID 10097), in which the anisotropy is not as severe but still quite important. In our analysis, we employed the atomic model with PDB ID 7VDF fitted to the densities.

### 3.2. FSC-3D calculation

3D-FSC calculations were carried out comparing post-processed densities against the maps simulated from the associated atomic models employing 20° cones and leaving all the remaining parameters set to default. The simulated maps were generated using EMAN2 [30] at the FSC global resolution. The employed masks were calculated using the simulated densities and binarized at threshold 0.01. A soft edge of 5 pixels was added to the masks.

### 3.3. New anisotropy method

As mentioned in the introduction, current approaches to evaluate the presence of anisotropic resolution in reconstructed maps are affected by the actual shape of the macromolecule. Thus, the resolution estimation through different orientations is convoluted by the actual shape of the macromolecule. Consequently, perfectly isotropic 3D reconstructions may be evaluated as anisotropic by current approaches (see Results section “Common map anisotropy metrics are affected by the shape of the specimen” and Figure 1). A good approach to evaluate the possibility of having preferred orientations and thus anisotropic reconstructions is to compute the distribution of aligned particles along the different possible orientations. However, this information, which is only an estimation, is not usually provided by the authors, so it is not possible to calculate it *a posteriori* if only the maps are made public.

To robustly evaluate the presence of preferred orientations in cryo-EM reconstructions, we have developed an approach that is not affected by the shape of the reconstructed macromolecule. Our proposed method uses the noise region outside the expected macromolecular density to estimate the noise power directional distribution. Our rationale is that the noise present in the solvent mask should show isotropic power through different directions for reconstructions not affected by preferential orientations, irrespective of the shape of the macromolecule. It is well-known that popular Bayesian-based 3D reconstruction methods work in ‘global alignment mode’, in which each particle contributes to all possible orientations in the 3D reconstruction albeit with different weights or likelihoods. However, typical 3D refinement methods achieving high resolution structures such as Relion [19,20] or cryoSPARC [31] switch to ‘local alignment searches’ after the global alignment phase generates a map with sufficient resolution. At this stage, achieving further improvements requires finer-grain angular resolution, which would make the computational cost of global alignment prohibitive; thus, local searches are used instead. In local searches, the weights for particles that deviate much from the most probable orientation, according to the previous iteration, are set to zero, and only those that are near the most probable orientation are indeed considered. In this case, the weighted average for each orientation is obtained only for particles that were accurately aligned at each orientation in the previous iteration. Thus, for 3D maps reconstructed after local angular searches, the obtained map weighted average (𝑉_𝑗_) at orientation j is given by,

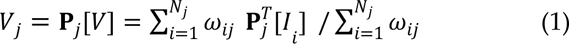

with 𝜔_𝑖𝑗_ the corresponding weight or likelihood for particle 𝑖 at orientation *j*, 𝐼_𝑖_ the Fourier Transform of the particle 𝑖, 𝑁_𝑗_ the number of particles locally aligned and weighted averaged at orientation *j* and operators 𝐏_𝑗_ and 𝐏^𝑇^ that extracts a central slice out of the 3D Fourier transform at orientation j and that places the 2D Fourier transform of an image back into the 3D transform respectively. By analysing the noise of the particles and not the macromolecule signal, (this can be achieved by applying a spherical soft mask to filter out the macromolecule after obtaining the orientation of the particles) we can study the anisotropy of the noise, which is not affected by the actual shape of the macromolecule. From Eq. (1), we can see that there is a direct relation between the obtained noise power at orientation j and the particle population at this orientation. The higher the 𝑁_𝑗_, the more effective the noise cancellation and, the lower the 𝑉_𝑗_ value will be.

As depicted in Figure 10, the noise power orientation distribution can be computed from the Fourier transform of the reconstructed map once it is previously filtered with a spherical soft mask. This power orientation distribution can provide direct insights into the particle orientation distribution without the need of analysing the set of particles used for the reconstruction. In practise, we estimate the noise power (NP) orientation distribution from the reconstructed map between two given resolutions, typically among 10 Å (𝑟_𝑚𝑖𝑛_) and the Nyquist resolution (𝑟_𝑚𝑎𝑥_) by averaging the noise power for all resolutions r between 𝑟_𝑚𝑖𝑛_ and 𝑟_𝑚𝑎𝑥_ at orientation j, and normalizing this quantity by its maximum noise power response as,

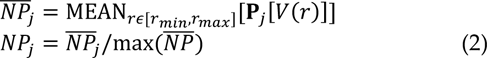

**Figure 10.**
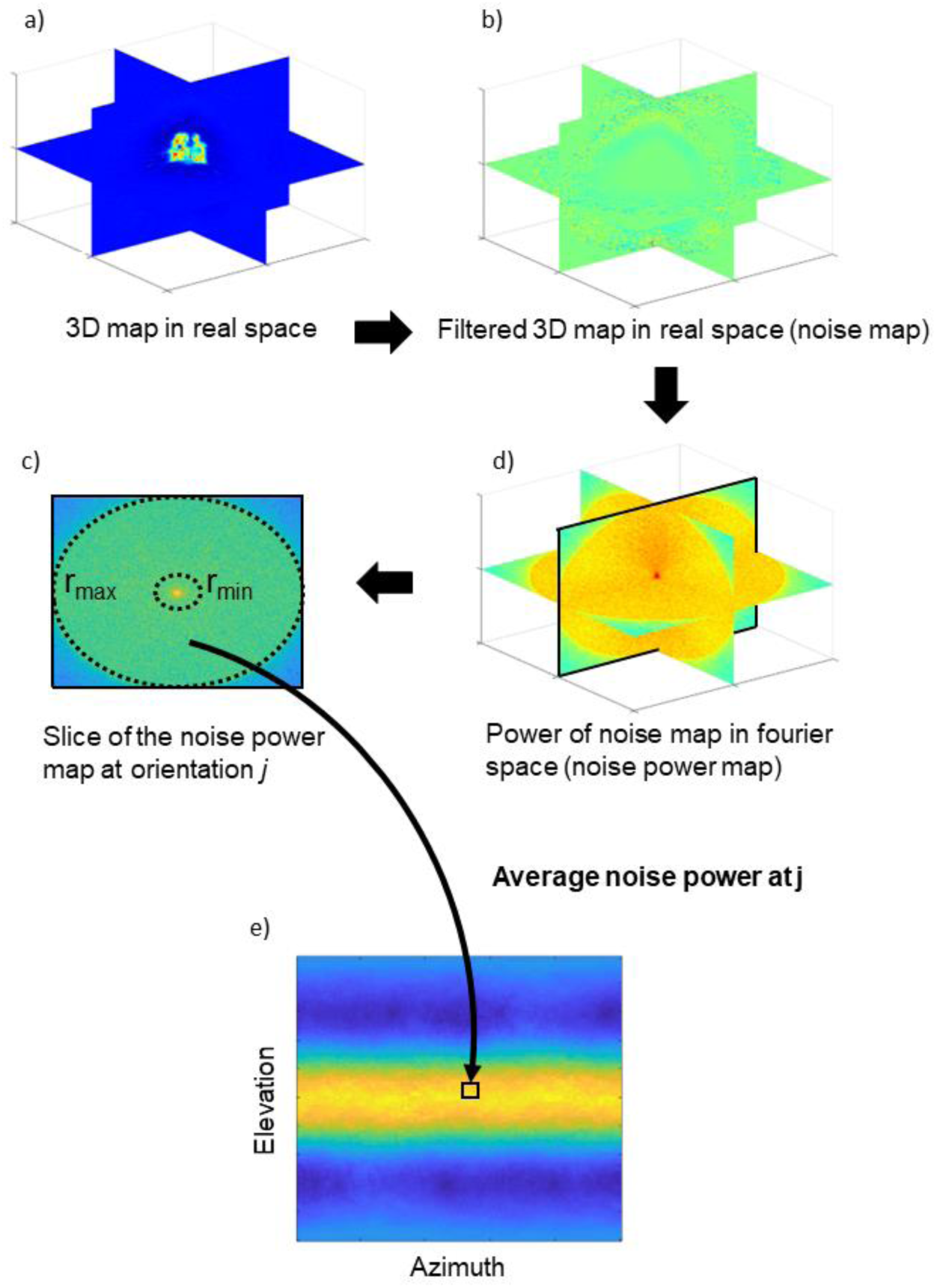
Workflow of the proposed anisotropy estimation method. The real space reconstructed 3D map (a) is filtered with a 3D spherical soft mask to filter out the macromolecular signal leaving only the noise (b). The filtered map is then transformed to the Fourier space and its 3D power map is calculated computing the square modulus of each Fourier component (d). Then, for each central slice of the power map the average noise power is calculated between given r_min_ and r_max_ resolutions (c). The central slice at orientation j is used to fill up the value of the noise power orientation distribution at this orientation j. As a result, the noise power directional distribution is obtained (e), the values in this distribution are normalized as shown in Eq. (2). Note that in (d) we show the logarithm of the power map for visualization purposes.

Special attention should be given to empty orientations without noise particles assigned. Note that for these cases, 𝑉_𝑗_ typically will not be zero despite 𝑁_𝑗_ being zero. This occurs because popular gridding-based direct Fourier reconstruction methods use a weighting approach where Fourier particle slices contribute to 3D map Fourier samples that are distant from them [19,32,33]. Therefore, Eq. (1) and (2) are applicable even for empty orientations, as the map Fourier components at these orientations will usually be interpolated from neighbour particle Fourier slices. However, this rationale does not apply for cases where large map Fourier regions are completely empty so they cannot be interpolated from neighbour slices. Nevertheless, these problematic cases can be easily detected by our approach as they will correspond to map orientations with zero or very close to zero noise power (𝑉_𝑗_).

The proposed anisotropic method should be used for 3D maps obtained by single particle cryo-EM, which have been computed with the typical cryo-EM pipeline using popular software packages such as Relion and/or cryoSPARC, that employ gridding-based direct Fourier reconstruction methods [19,32,33], providing robustness to potential empty regions in Fourier space. We also assume that the map was not been postprocessed by any approach, such as sharpening [15,22–25], map enhancement [21,34] or density modification [35] methods. Additionally, we also presume that the map was not computed using complicated processing schemas as for example sub-particle masked refinement or density subtraction, as our assumptions about noise power may not hold in these scenarios.

However, this approach might still be useful for 3D reconstructions obtained by other reconstruction methods as projection matching [36] or even subtomogram averaging [37] in electron cryotomography, providing the reconstructed maps approximately sample well the map Fourier 3D grid without leaving major gaps. Therefore, this method should not be applied to maps reconstructed from tomography tilt series, as they are affected by the missing wedge issue, and they will show important empty regions with no information in the Fourier map. Even in such cases, problematic empty Fourier regions with no information can be easily detected in the 2D map obtained by our approach as it will show zero or close to zero noise power (𝑉_𝑗_) for the corresponding orientations. We do not anticipate that the proposed method will have problems to analyze reconstructions affected by overfitting or “ghost” densities coming from close particles inside the particle box as far as these effects are isotropic and do not show any preferred directions.

Our approach first defines a spherical soft mask to filter out the macromolecular signal, leaving only noise. Then, to analyse the anisotropy of the noise, the Fourier Transform of the masked map is computed and slices at different orientations are extracted at the elevation and azimuth Euler’s angles of the unit vectors perpendicular to the slices. In other words, an elevation angle given by 1.5 rad (or −1.5 rad) corresponds to a top view projection, while an elevation of 0 rad corresponds to a side view projection. For each extracted slice, the average noise power is computed for a defined resolution range between 10 Å and the corresponding Nyquist resolution. Finally, the highest computed noise power along all computed directions is used to normalize a 2D heatmap, that represents the normalized noise power for different elevation and azimuth Euler’s angles.

Thus, the proposed approach can provide effectively *ex post facto* information of the distribution of particles along projection direction, without requiring any angular assignment data. Additionally, it is worth noting that particle orientation distributions are often calculated without considering the particle’s weight distributions, using only the orientation with highest weight for each particle. However, cryo-EM reconstructions are not usually computed in this way since a particle may contribute to several orientations. As explained, our proposed approach can evaluate the presence of anisotropic particle distributions taking into consideration the particlés weight distributions used in the reconstruction, and it is insensitive to the shape of the macromolecule.

### 3.4. Map post-processing

Five different map postprocessing algorithms were employed: DeepEMhancer [21], LocScale2 [22,23], LocSpiral [24], Phenix anisotropic sharpening [15], and LocalDeblur [25]. DeepEMhancer was executed with default parameters for all cases. Phenix phenix.local_aniso_sharpen was executed with local_sharpen=True anisotropic_sharpen=True. LocScale2 was executed with default parameters using the atomic model, the global FSC resolution and the half maps as inputs. LocSpiral was executed with a threshold of 0.9 and a bandwidth manually picked at 4.5. LocSpiral and LocScale2 were executed using the same masks that were used later for 3D-FSC estimation. The model-map LocScale2 pipeline was executed with default parameters. Local resolution maps required for LocalDeblur were computed using Monores [38] using as inputs the half-maps and the same mask that in LocSpiral and LocScale2 executions.

### 3.5. Map visualisation

Figures 7 and 8 that display densities around atomic model locations were generated using UCSF ChimeraX version 1.6.1 [39,40] and ISOLDE version 1.6.0 [41]. Atomic models (PDB entries 6UJA and 7VDF) were rigidbody fitted into the corresponding maps (EMD-20794 and EMD-8731, respectively) and subsequently inspected exhaustively using the residue stepper command from ISOLDE. Contour levels were chosen to initially match the ones used for the map overview figures (see above) and were adjusted when necessary to best visualize local features (all contour levels are indicated in the figures). No refinement other than a global rigid-body fit into the map was carried out on the atomic models, since they were only used for navigation and illustration.

Supplementary Figures S7 and S8 displaying the whole maps were generated using UCSF ChimeraX version 1.2 [39,40]. The threshold values employed in Figure S7 were 0.672, 0.098, 4.730, 0.864, 0.123, 0.820 for the Raw, DeepEMhancer, LocSpiral, LocalDeblur, LocScale and Phenix sharpened maps and 0.0281, 0.223, 6.03, 0.0653, 0.045 and 0.0744 in Figure S8. In Figure S8, the displayed maps were masked using a mask derived from 6UJA for comparison ease. The mask was generated by simulating a map from an atomic model using ChimeraX molmap at 4 Å resolution following by a gaussian smoothing with sigma 2 Å and volume falloff with 10 iterations. As a mask threshold, the percentile 10 was selected.

## 4. Conclusions

The preferred orientations that some specimens tend to exhibit remain as a major challenge in cryo-EM. Preferred orientations are the main cause of the resolution anisotropy and due to their relevance, several directional resolution methods have been proposed as proxy metrics for the preferential orientation problem. Some of these metrics have been used to evaluate newly proposed experimental approaches aimed to minimise the problem of preferred orientations, but all of them present an important limitation: they are sensitive to the shape of the specimen and consequently, inappropriate for large scale experiments comparing different samples. Our newly proposed method for preferred orientation analysis has been designed to explicitly account for this short-coming.

Although this limitation may not be considered an important issue, if new approaches to tackle the problem of preferred orientations are to be proposed, robust metrics for its estimation are a requirement to assess their effectiveness across multi-specimen benchmarks. Even though an estimated distribution of single particle orientations can be obtained from widely used cryo-EM software, this information is not always included in published papers. In addition, supporting cryo-EM movies/micrographs/particles are not always deposited in EMPIAR [42] database, while raw cryo-EM maps are always required to be deposited in the EMDB database, which explains the popularity of other map anisotropy methods such as 3D-FSC. Our proposed method does not require the particle projection distribution to evaluate the potential presence of anisotropy and only requires the raw cryo-EM map reconstruction, thus, it can be used to evaluate controversial maps *ex post facto* for which we do not have particle information available, or even cases where authors include this information but there are doubts about its accuracy.

While currently there is only one computational approach explicitly designed to tackle the problem of map anisotropy, in this manuscript we have shown that some of the most recently published map post-processing algorithms are also capable of attenuating map anisotropy. According to our results, these algorithms can enhance the visualisation of maps affected with moderate amounts of anisotropy with negligible additional computing cost, making them potentially useful for projects in which the preferred orientations problem is not severe enough to prevent the reconstruction process, but it is strong enough to affect the quality and interpretability of the map.

On the other hand, we must be very careful when trying to interpret maps affected by anisotropy that have been enhanced by postprocessing algorithms. As mentioned in the introduction, map anisotropy is a consequence of an experimental deficit. Thus, attempting to computationally correct map anisotropy by introducing *a priori* information presents significant risks. Note that currently there is no possible way to validate whether this inpainting of unobserved data accurately represents the actual underlying structure, or not. Consequently, although enhanced postprocessed maps can be used to guide the tracing of the atomic model, the experimental map should always be used to refine the atomic model and to confirm or not the presence of the features suggested by the postprocessed maps. We have found the combined use of both the experimental and postprocessed maps in interactive model building and refinement tools such as ISOLDE [41] to be highly advantageous. This approach consists of opening both maps simultaneously but exclusively utilizing the experimental map to drive the molecular dynamics flexible fitting [43] procedure, with the postprocessed map used solely as a visual guide. Very often, for protein residues around which the experimental map is difficult to interpret, a postprocessed map suggests a location for side-chain atoms where the combined pull of the experimental map and molecular dynamics force field result in a stable conformation.

To the best of our knowledge, this is the first study to show that existing algorithms can significantly improve map visualizations affected by preferred orientations. As we look to the future, we anticipate that this avenue will invite more extensive research, not just to harness its promising potential (e.g., [44]), but also to rigorously evaluate and navigate the accompanying risks.

## Code availability

Code to compute the map anisotropy and to generate the analysis figures can be found at https://github.com/1aviervargas/detectAnisotropy. Postprocessed maps can be download at https://zenodo.org/record/8399272.

## Acknowledgments

We would like to thank our colleagues for interesting discussions and feedback. Ruben Sanchez Garcia is a post-doctoral fellow funded via an Astex Pharmaceuticals Sustaining Innovation Post-Doctoral Fellowship. We acknowledge financial support from the Spanish Ministerio de Ciencia e Innovación, Grant PID2022-137548OB-I00 funded by MCIN/AEI/10.13039/501100011033/ and by ERDF A way of making Europe.

## Authors’ contributions

Conceptualization, Ruben Sánchez-García and Javier Vargas; Methodology, Ruben Sánchez-García and Javier Vargas; Software, Ruben Sánchez-García and Javier Vargas; Validation, Ruben Sánchez-García, Guillaume Gaullier and Javier Vargas; Investigation, Ruben Sánchez-García; Data curation, Guillaume Gaullier; Writing – original draft, Ruben Sánchez-García and Javier Vargas; Writing – review & editing, Ruben Sánchez-García, Jose Manuel Cuadra-Troncoso and Javier Vargas; Funding acquisition, Javier Vargas.

## Supporting information

**Figure S1.**
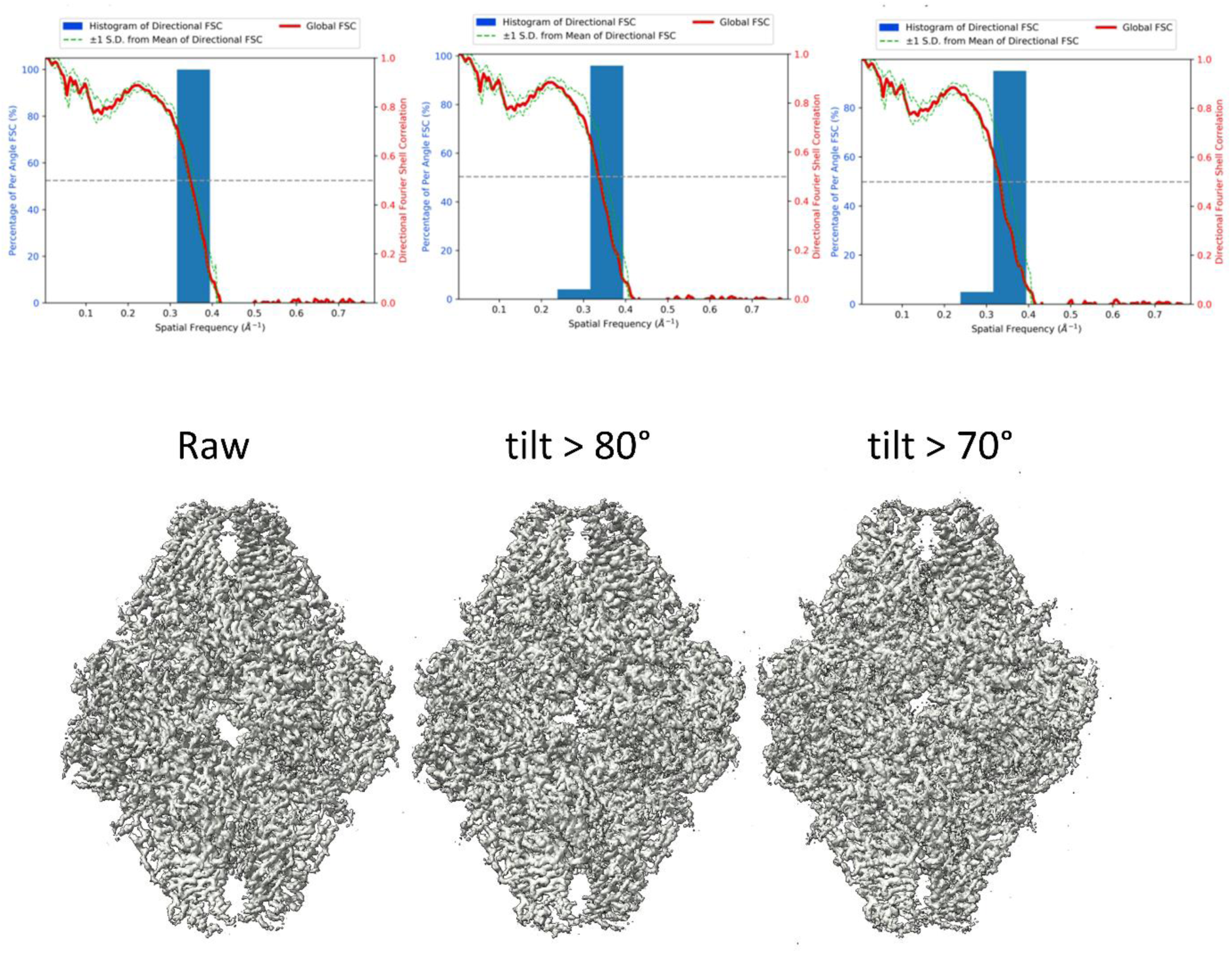
Bottom, cryo-EM maps for the β-galactosidase reconstructed from the EMPIAR-10061 dataset, using particles from all the projection directions (Left), and removing 95% of the particles with tilt angle >80° (Middle) and >70° (Right). Top, 3D-FSC displaying the global FSC resolution (red) and standard deviation of the directional resolution over 20-degree cones (green) as well as the histogram of the per angle FSC for the raw β-galactosidase map (Left), and the maps reconstructed with 95% of the particles with tilt angle >80° (Middle) and >70° (Right) removed.

**Figure S2.**
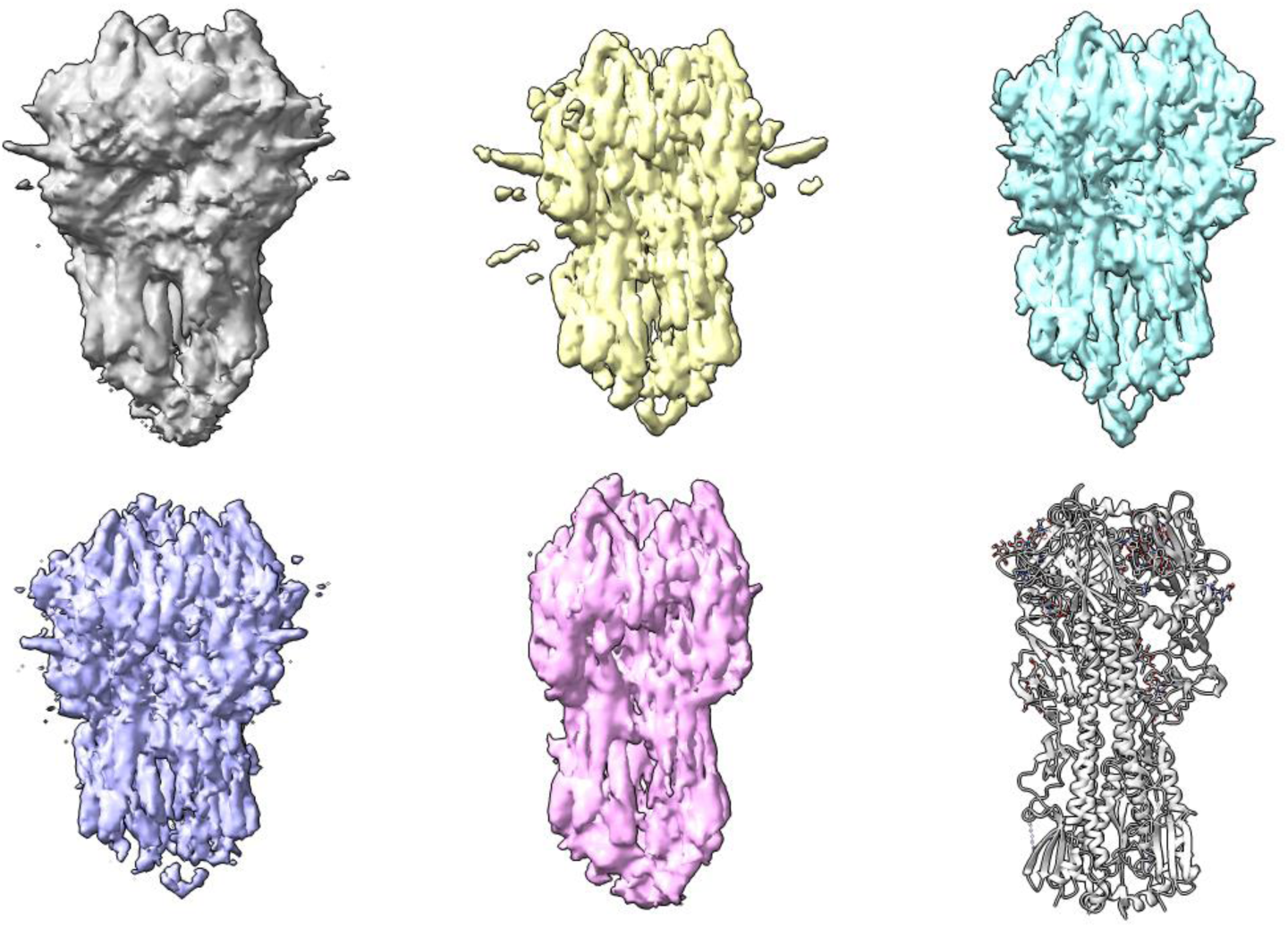
Hemagglutinin trimer map reconstructed from untilted micrographs (grey), and the results of applying DeepEMhancer (yellow, [29]), LocSpiral (cyan, [27]), LocalDeblur (purple, [26]), and LocScale2 (pink, [25,28]) to the reconstructed map. The atomic model used for evaluation is depicted in pale grey.

**Figure S3.**
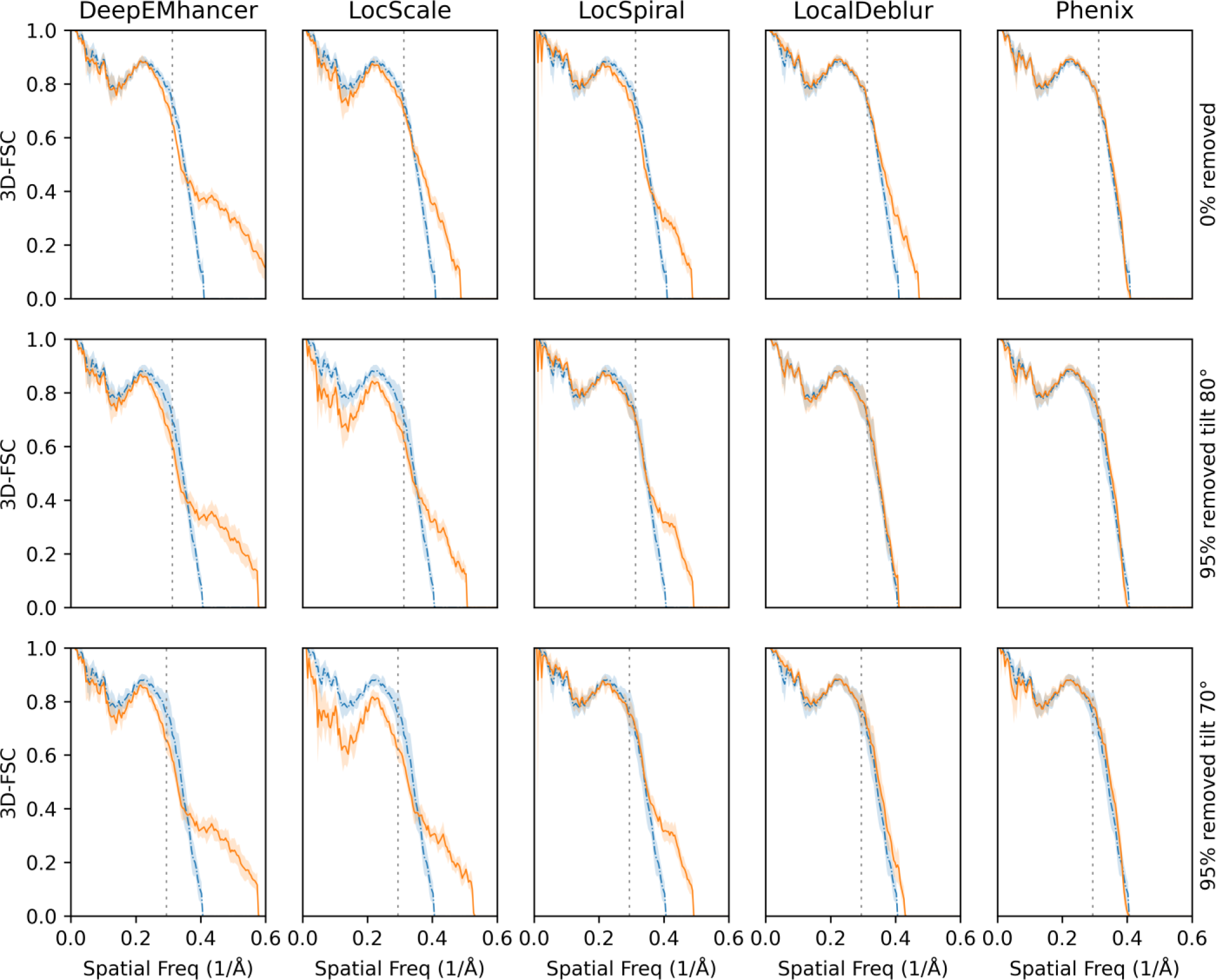
Alternative representation of the per-cone map to model FSC (mean, solid lines, standard deviation, shadow) calculated with 3D-FSC for three different versions of the β-galactosidase complex: reconstructed with all particles (Top) and reconstructed with 95% of the particles with tilt angle >80° (Middle) and >70° (Bottom) removed. The blue line represents the reconstructed map whereas the orange lines represent the maps obtained when DeepEMhancer [29], LocScale2 [25,28], LocSpiral [27], Phenix [14] and LocalDeblur [26] are employed on the reconstructed map. The FSC global resolution is displayed as a dotted vertical line.

**Figure S4.**
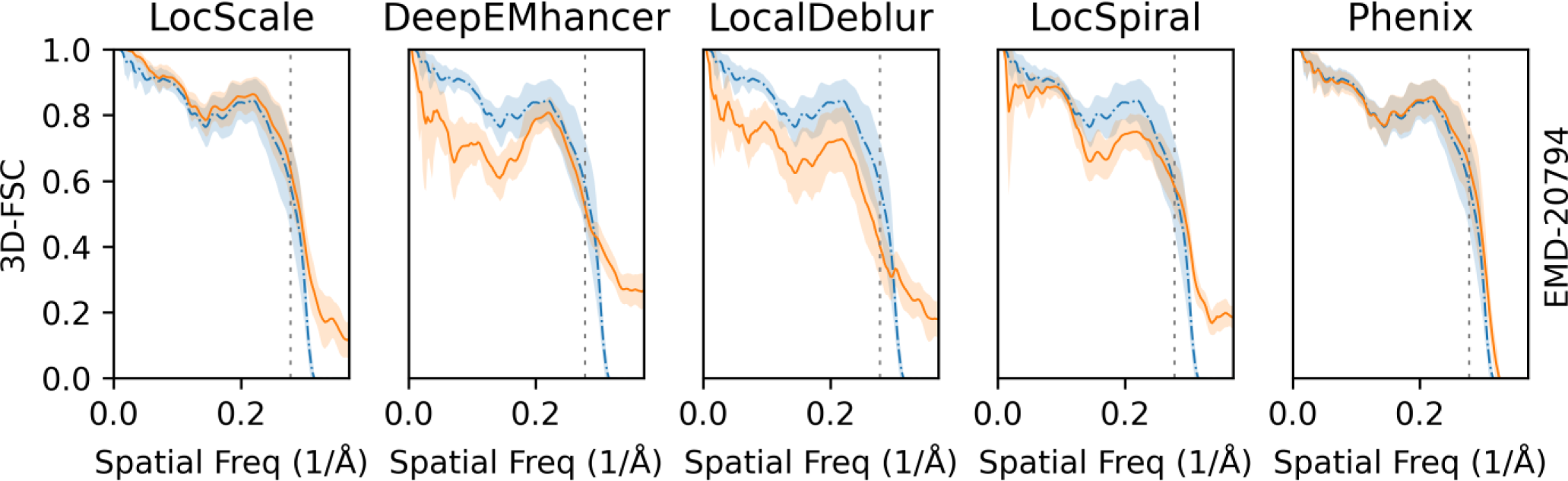
Alternative representation of the per-cone map to model FSC (mean, solid lines, standard deviation, shadow) for the EMD-20794. The blue line represents the reconstructed map whereas the orange lines represent the standard deviation for the maps obtained when DeepEMhancer [29], LocScale2 [25,28], LocSpiral [27], Phenix [14] and LocalDeblur [26] are employed on the reconstructed map. The FSC global resolution is displayed as a dotted vertical line.

**Figure S5.**
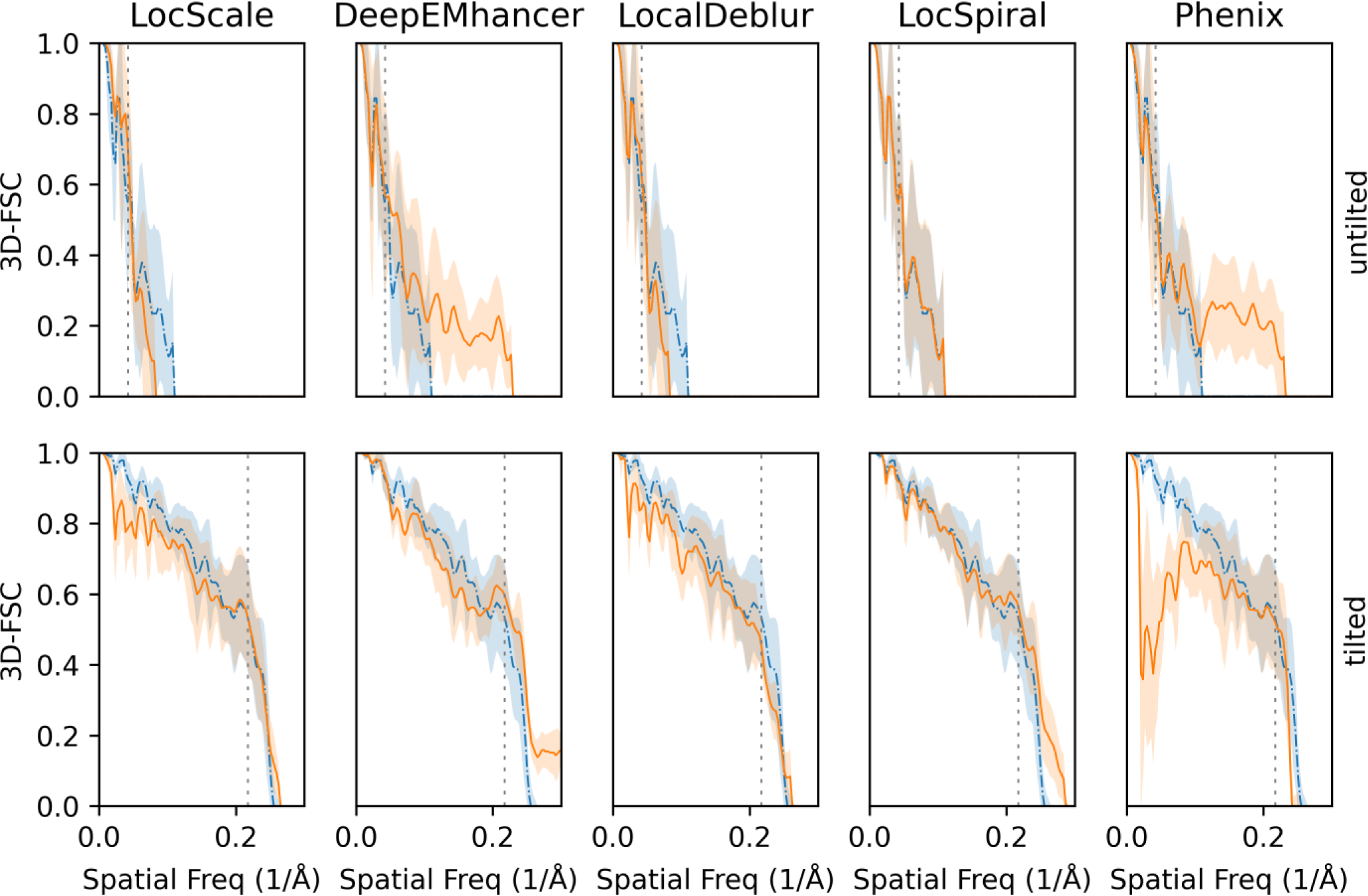
Alternative representation of the per-cone map to model FSC (mean, solid lines, standard deviation, shadow) calculated with 3D-FSC for two different versions of the influenza hemagglutinin trimer: reconstructed with untilted micrographs (severe anisotropy problems) and reconstructed with micrographs tilted 40°. The blue line represents the reconstructed map whereas the orange lines represent the maps obtained when DeepEMhancer [29], LocScale2 [25,28], LocSpiral [27], Phenix [14] and LocalDeblur [26] are employed on the reconstructed map. The FSC global resolution is displayed as a dotted vertical line.

**Figure S6.**
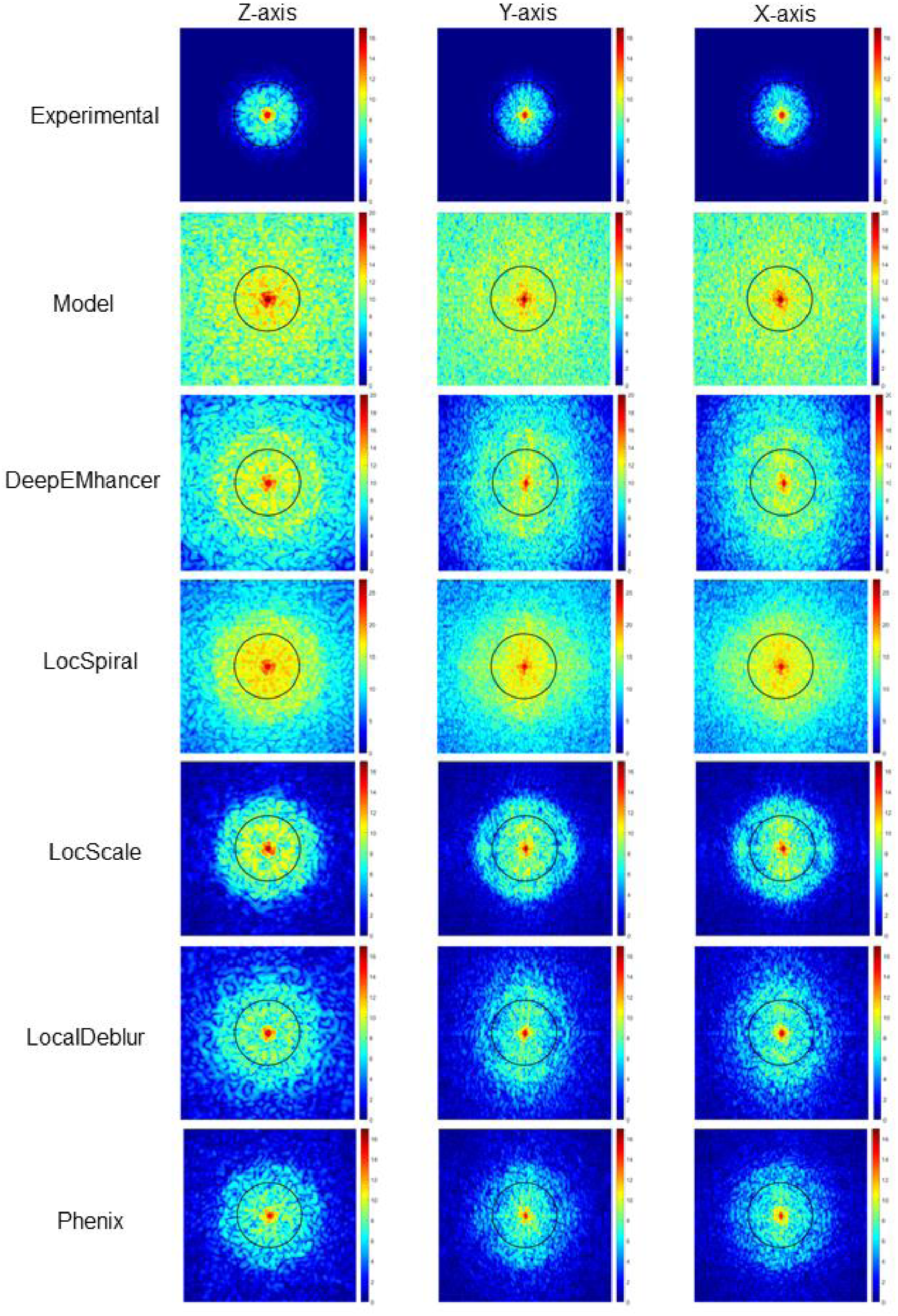
Logarithm of Fourier amplitudes for the tilted influenza hemagglutinin trimer reconstructions at slices *z* = 0, *y* = 0 and *x* = 0 after running the different postprocessing approaches. The first and second rows correspond to the experimental map without any postprocessing (first row) and the map simulated from the atomic model (second row). The black circle in the figures shows a resolution corresponding to 3.5 Å.

**Figure S7.**
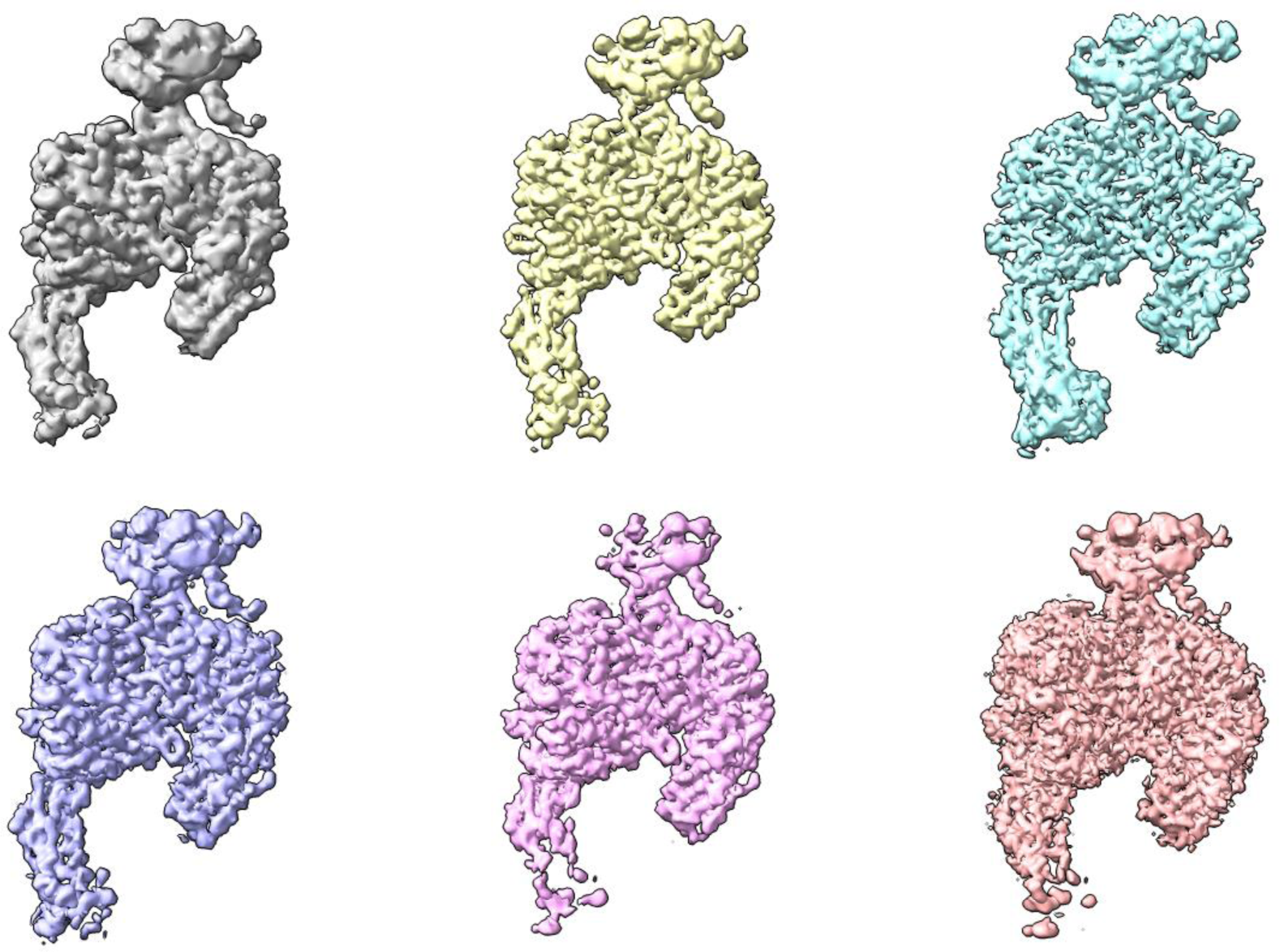
Reconstructed map (grey) for the EMD-20794 and the results of applying DeepEMhancer (yellow, [29]), LocSpiral (cyan, [27]), LocalDeblur (purple, [26]), LocScale2 (pink, [25,28]) and Phenix anisotropic sharpening (salmon, [14]) to the reconstructed map.

**Figure S8.**
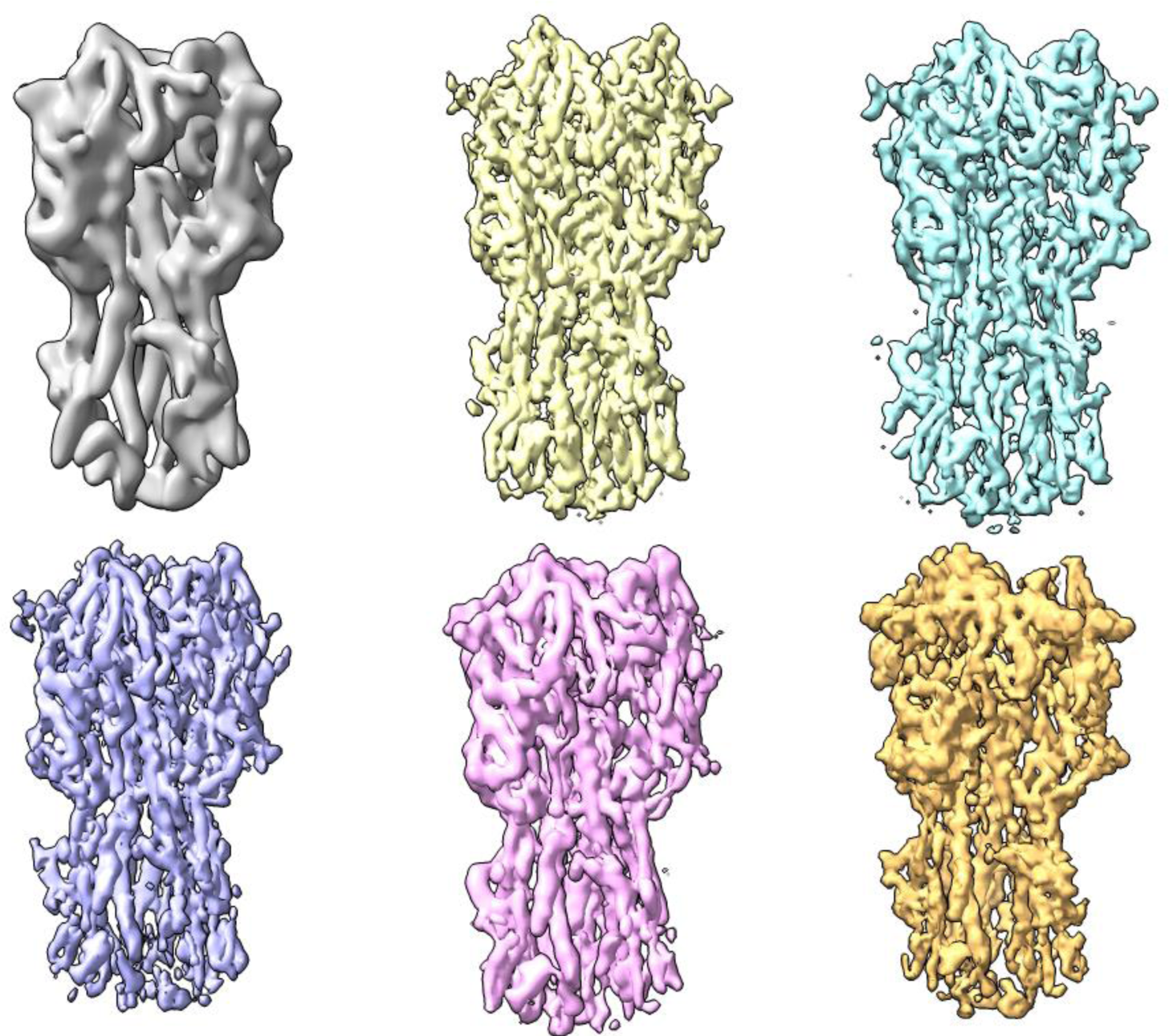
Influenza hemagglutinin trimer map reconstructed from tilted micrographs (grey), and the results of applying DeepEMhancer (yellow, [29]), LocSpiral (cyan, [27]), LocalDeblur (purple, [26]), LocScale2 (pink, [25,28]) and Phenix anisotropic sharpening (salmon, [14]) to the reconstructed map. The atomic model used for evaluation is depicted in pale grey.

## References

1. Kühlbrandt, W. The Resolution Revolution. Science 2014, 343, 1443–1444, doi:10.1126/science.1251652

2. D’Imprima, E.; Kühlbrandt, W. Current Limitations to High-Resolution Structure Determination by Single-Particle CryoEM. Q Rev Biophys 2021, 54, e4, doi:10.1017/S0033583521000020.

3. Naydenova, K.; Russo, C.J. Measuring the Effects of Particle Orientation to Improve the Efficiency of Electron Cryomicroscopy. Nature Communications 2017 8:1 2017, 8, 1–5, doi:10.1038/s41467-017-00782-3.

4. Noble, A.J.; Dandey, V.P.; Wei, H.; Brasch, J.; Chase, J.; Acharya, P.; Tan, Y.Z.; Zhang, Z.; Kim, L.Y.; Scapin, G.;, et al. Routine Single Particle CryoEM Sample and Grid Characterization by Tomography. Elife 2018, 7, doi:10.7554/ELIFE.34257.

5. Chua, E.Y.D.; Mendez, J.H.; Rapp, M.; Ilca, S.L.; Tan, Y.Z.; Maruthi, K.; Kuang, H.; Zimanyi, C.M.; Cheng, A.; Eng, E.T.;, et al. Better, Faster, Cheaper: Recent Advances in Cryo–Electron Microscopy. 10.1146/annurev-biochem-032620-110705 2022, 91, 1–32, doi:10.1146/ANNUREV-BIOCHEM-032620-110705.

6. Zi Tan, Y.; Baldwin, P.R.; Davis, J.H.; Williamson, J.R.; Potter, C.S.; Carragher, B.; Lyumkis, D. Addressing Preferred Specimen Orientation in Single-Particle Cryo-EMthrough Tilting. Nat Methods 2017, 14, 793–796, doi:10.1038/nmeth.4347.

7. Sorzano, C.O.S.; Semchonok, D.; Lin, S.C.; Lo, Y.C.; Vilas, J.L.; Jiménez-Moreno, A.; Gragera, M.; Vacca, S.; Maluenda, D.; Martínez, M.;, et al. Algorithmic Robustness to Preferred Orientations in Single Particle Analysis by CryoEM. J Struct Biol 2021, 213, 107695, doi:10.1016/J.JSB.2020.107695.

8. Vilas, J.L.; Tagare, H.D.; Vargas, J.; Carazo, J.M.; Sorzano, C.O.S. Measuring Local-Directional Resolution and Local Anisotropy in Cryo-EM Maps. Nature Communications 2020 11:1 2020, 11, 1–7, doi:10.1038/s41467-019-13742-w.

9. Vilas, J.; Tagare, H.D. Three New Measures of Anisotropy of Cryo-EM Maps Three New Measures of Anisotropy of Cryo-EM Maps. 2022, 0–14, doi:10.21203/RS.3.RS-1585291/V1.

10. Glaeser, R.M.; Han, B.-G. Opinion: Hazards Faced by Macromolecules When Confined to Thin Aqueous Films. Biophys Rep 2017, 3, 1–7, doi:10.1007/S41048-016-0026-3.

11. Russo, C.J.; Passmore, L.A. Controlling Protein Adsorption on Graphene for Cryo-EM Using Low-Energy Hydrogen Plasmas. Nature Methods 2014 11:6 2014, 11, 649–652, doi:10.1038/nmeth.2931.

12. Han, B.G.; Walton, R.W.; Song, A.; Hwu, P.; Stubbs, M.T.; Yannone, S.M.; Arbeláez, P.; Dong, M.; Glaeser, R.M. Electron Microscopy of Biotinylated Protein Complexes Bound to Streptavidin Monolayer Crystals. J Struct Biol 2012, 180, 249–253, doi:10.1016/J.JSB.2012.04.025.

13. Wang, F.; Liu, Y.; Yu, Z.; Li, S.; Feng, S.; Cheng, Y.; Agard, D.A. General and Robust Covalently Linked Graphene Oxide Affinity Grids for High-Resolution Cryo-EM. Proc Natl Acad Sci U S A 2020, 117, 24269– 24273, doi:10.1073/PNAS.2009707117/SUPPL_FILE/PNAS.2009707117.SM02.MP4.

14. Noble, A.J.; Wei, H.; Dandey, V.P.; Zhang, Z.; Tan, Y.Z.; Potter, C.S.; Carragher, B. Reducing Effects of Particle Adsorption to the Air–Water Interface in Cryo-EM. Nature Methods 2018 15:10 2018, 15, 793–795, doi:10.1038/s41592-018-0139-3.

15. Terwilliger, T.C.; Sobolev, O. V.; Afonine, P. V.; Adams, P.D. Automated Map Sharpening by Maximization of Detail and Connectivity. Acta Crystallogr D Struct Biol 2018, 74, 545–559, doi:10.1107/S2059798318004655.

16. Peter, M.F.; Ruland, J.A.; Depping, P.; Schneberger, N.; Severi, E.; Moecking, J.; Gatterdam, K.; Tindall, S.; Durand, A.; Heinz, V.;, et al. Structural and Mechanistic Analysis of a Tripartite ATP-Independent Periplasmic TRAP Transporter. Nature Communications 2022 13:1 2022, 13, 1–15, doi:10.1038/s41467-022-31907-y.

17. Melo, A.A.; Sprink, T.; Noel, J.K.; Vázquez-Sarandeses, E.; van Hoorn, C.; Mohd, S.; Loerke, J.; Spahn, C.M.T.; Daumke, O. Cryo-Electron Tomography Reveals Structural Insights into the Membrane Remodeling Mode of Dynamin-like EHD Filaments. Nature Communications 2022 13:1 2022, 13, 1–13, doi:10.1038/s41467-022-35164-x.

18. Rosenthal, P.B.; Henderson, R. Optimal Determination of Particle Orientation, Absolute Hand, and Contrast Loss in Single-Particle Electron Cryomicroscopy. J Mol Biol 2003, 333, 721–745, doi:10.1016/j.jmb.2003.07.013.

19. Scheres, S.H.W. RELION: Implementation of a Bayesian Approach to Cryo-EM Structure Determination. J Struct Biol 2012, 180, 519–530, doi:10.1016/j.jsb.2012.09.006.

20. Zivanov, J.; Nakane, T.; Forsberg, B.O.; Kimanius, D.; Hagen, W.J.H.; Lindahl, E.; Scheres, S.H.W. New Tools for Automated High-Resolution Cryo-EM Structure Determination in RELION-3. Elife 2018, 7, e42166 doi:10.7554/eLife.42166.

21. Sanchez-Garcia, R.; Gomez-Blanco, J.; Cuervo, A.; Carazo, J.M.; Sorzano, C.O.S.; Vargas, J. DeepEMhancer: A Deep Learning Solution for Cryo-EM Volume Post-Processing. Commun Biol 2021, 4, 1–8.

22. Jakobi, A.J.; Wilmanns, M.; Sachse, C. Model-Based Local Density Sharpening of Cryo-EM Maps. Elife 2017, 6.

23. Bharadwaj, A.; Jakobi, A.J. Electron Scattering Properties of Biological Macromolecules and Their Use for Cryo-EM Map Sharpening. Faraday Discuss 2022, doi:10.1039/D2FD00078D.

24. Kaur, S.; Gomez-Blanco, J.; Khalifa, A.A.Z.; Adinarayanan, S.; Sanchez-Garcia, R.; Wrapp, D.; McLellan, J.S.; Bui, K.H.; Vargas, J. Local Computational Methods to Improve the Interpretability and Analysis of Cryo-EM Maps. Nat Commun 2021, 12, 1240, doi:10.1038/s41467-021-21509-5.

25. Ramírez-Aportela, E.; Vilas, J.L.; Glukhova, A.; Melero, R.; Conesa, P.; Martínez, M.; Maluenda, D.; Mota, J.; Jiménez, A.; Vargas, J.;, et al. Automatic Local Resolution-Based Sharpening of Cryo-EM Maps. Bioinformatics 2020, 36, 765–772, doi:10.1093/bioinformatics/btz671.

26. Bartesaghi, A.; Merk, A.; Banerjee, S.; Matthies, D.; Wu, X.; Milne, J.L.S.; Subramaniam, S. 2.2 Å Resolution Cryo-EM Structure of β-Galactosidase in Complex with a Cell-Permeant Inhibitor. Science 2015, 348, 1147– 1151, doi:10.1126/SCIENCE.AAB1576.

27. Campbell, M.G.; Cormier, A.; Ito, S.; Seed, R.I.; Bondesson, A.J.; Lou, J.; Marks, J.D.; Baron, J.L.; Cheng, Y.; Nishimura, S.L. Cryo-EM Reveals Integrin-Mediated TGF-β Activation without Release from Latent TGF-β. Cell 2020, 180, 490–501.e16, doi:10.1016/j.cell.2019.12.030.

28. Oli B. Clarke, @OliBClarke, 2020, Twitter. https://twitter.com/OliBClarke/status/1301985078231232512.

29. Campbell, M.G.; Cormier, A.; Ito, S.; Seed, R.I.; Bondesson, A.J.; Lou, J.; Marks, J.D.; Baron, J.L.; Cheng, Y.; Nishimura, S.L. Cryo-EM Reveals Integrin-Mediated TGF-β Activation without Release from Latent TGF-β. Cell 2020, 180, 490–501.e16, doi:10.1016/j.cell.2019.12.030.

30. Tang, G.; Peng, L.; Baldwin, P.R.; Mann, D.S.; Jiang, W.; Rees, I.; Ludtke, S.J. EMAN2: An Extensible Image Processing Suite for Electron Microscopy. J Struct Biol 2007, 157, 38–46, doi:10.1016/j.jsb.2006.05.009.

31. Punjani, A.; Rubinstein, J.L.; Fleet, D.J.; Brubaker, M.A. CryoSPARC: Algorithms for Rapid Unsupervised Cryo-EM Structure Determination. Nature Methods 2017 14:3 2017, 14, 290–296, doi:10.1038/nmeth.4169.

32. Pipe, J.G.; Menon, P. Sampling Density Compensation in MRI: Rationale and an Iterative Numerical Solution. Magn Reson Med 1999, 41, 179–186, doi:10.1002/(SICI)1522-2594(199901)41:1.

33. Střelák, D.; Sorzano, C.Ó.S.; Carazo, J.M.; Filipovič, J. A GPU Acceleration of 3-D Fourier Reconstruction in Cryo-EM. The International Journal of High Performance Computing Applications 2019, 33, 948–959, doi:10.1177/1094342019832958.

34. Vargas, J.; Gómez-Pedrero, J.A.; Quiroga, J.A.; Quiroga, J.A.; Alonso, J.; Alonso, J. Enhancement of Cryo-EM Maps by a Multiscale Tubular Filter. Optics Express, Vol. 30, Issue 3, pp.4515–4527 2022, 30, 4515–4527, doi:10.1364/OE.444675.

35. Terwilliger, T.C.; Ludtke, S.J.; Read, R.J.; Adams, P.D.; Afonine, P. V. Improvement of Cryo-EM Maps by Density Modification. Nature Methods 2020 17:9 2020, 17, 923–927, doi:10.1038/s41592-020-0914-9.

36. Penczek, P.A.; Grassucci, R.A.; Frank, J. The Ribosome at Improved Resolution: New Techniques for Merging and Orientation Refinement in 3D Cryo-Electron Microscopy of Biological Particles. Ultramicroscopy 1994, 53, 251–270, doi:10.1016/0304-3991(94)90038-8.

37. Wan, W.; Briggs, J.A.G. Cryo-Electron Tomography and Subtomogram Averaging. Methods Enzymol 2016, 579, 329–367, doi:10.1016/BS.MIE.2016.04.014.

38. Vilas, J.L.; Gómez-Blanco, J.; Conesa, P.; Melero, R.; Miguel de la Rosa-Trevín, J.; Otón, J.; Cuenca, J.; Marabini, R.; Carazo, J.M.; Vargas, J.;, et al. MonoRes: Automatic and Accurate Estimation of Local Resolution for Electron Microscopy Maps. Structure 2018, 26, 337–344.e4, doi:10.1016/j.str.2017.12.018.

39. Goddard, T.D.; Huang, C.C.; Meng, E.C.; Pettersen, E.F.; Couch, G.S.; Morris, J.H.; Ferrin, T.E. UCSF ChimeraX: Meeting Modern Challenges in Visualization and Analysis. Protein Sci 2018, 27, 14–25, doi:10.1002/PRO.3235.

40. Pettersen, E.F.; Goddard, T.D.; Huang, C.C.; Meng, E.C.; Couch, G.S.; Croll, T.I.; Morris, J.H.; Ferrin, T.E. UCSF ChimeraX: Structure Visualization for Researchers, Educators, and Developers. Protein Sci 2021, 30, 70–82, doi:10.1002/PRO.3943.

41. Croll, T.I. ISOLDE: A Physically Realistic Environment for Model Building into Low-Resolution Electron-Density Maps. Acta Crystallogr D Struct Biol 2018, 74, 519–530, doi:10.1107/S2059798318002425.

42. Iudin, A.; Korir, P.K.; Somasundharam, S.; Weyand, S.; Cattavitello, C.; Fonseca, N.; Salih, O.; Kleywegt, G.J.; Patwardhan, A. EMPIAR: The Electron Microscopy Public Image Archive. Nucleic Acids Res. 2023, 51, D1503–D1511, doi:10.1093/nar/gkac1062.

43. Trabuco, L.G.; Villa, E.; Schreiner, E.; Harrison, C.B.; Schulten, K. Molecular Dynamics Flexible Fitting: A Practical Guide to Combine Cryo-Electron Microscopy and X-ray Crystallography. Methods 2009, 49, 174– 180, doi:10.1016/j.ymeth.2009.04.005.

44. Liu, Y.-T.; Hu, J.; Zhou, Z.H. Resolving the Preferred Orientation Problem in CryoEM Reconstruction with Self-Supervised Deep Learning. Microscopy and Microanalysis 2023, 29, 1918–1919, doi:10.1093/micmic/ozad067.991.

